# 5’-*S*-(3-aminophenyl)-5’-thioadenosine, a novel chemoprotective agent for reducing toxic side effects of fluorouracil in treatment of MTAP-deficient cancers

**DOI:** 10.1101/2024.04.17.589975

**Authors:** Si Zhang, Hui Xue, Nelson K.Y. Wong, Thomas Doerksen, Fuqiang Ban, Shawn Aderson, Stanislav Volik, Yen-Yi Lin, Zhongye Dai, Ivica Bratanovic, Hongwei Cheng, Colin C. Collins, Artem Cherkasov, Jeremy Wulff, Yuzhuo Wang

**Affiliations:** The Vancouver Prostate Centre and Department of Urologic Sciences, University of British Columbia, Vancouver, BC, Canada; Department of Experimental Therapeutics, BC Cancer Research Institute, Vancouver, BC, Canada; Department of Chemistry, University of Victoria, Victoria, BC, Canada

**Keywords:** 5’-S-(3-aminophenyl)-5’-thioadenosine, fluorouracil, MTAP-deficient cancers, Nucleobase analogue, S-methyl-5’-thioadenosine

## Abstract

**Background:** Nucleobase analogue (NBA) drugs are effective chemotherapeutics, but their clinical use is limited by severe side effects. Compelling evidence suggests the use of *S*-methyl-5’-thioadenosine (MTA) can selectively reduce NBA toxicity on normal tissues while maintaining the efficacy of NBAs on methylthioadenosine phosphorylase (MTAP)-deficient cancers. However, we found that MTA induced hypothermia at its effective dose, limiting its translational potential. We intended to find an MTA analogue that can exert MTA function while minimize the undesired side effects of MTA. Thus, such an analogue can be used in combination with NBAs in selectively targeting MTAP-deficient cancers.

**Methods:** We screened a library of MTA analogues for the following criteria: 1) being substrates of MTAP; 2) selectively protection on MTAP-expressing cells from NBA toxicity using *MTAP*-isogenic cell lines; 3) ability to protect the host from NBA toxicity without hypothermic effect; and 4) lack of interference on the tumor-suppressive effect of NBA in mice bearing MTAP-deficient tumors.

**Results:** We identified 5’-*S*-(3-aminophenyl)-5’-thioadenosine (m-APTA) that did not induce hypothermia at the effective doses. We demonstrated that m-APTA could be converted to adenine by MTAP. Consequently, m-APTA selectively protected mouse hosts from 5-FU-induced toxicity (i.e. anemia); yet it did not interfere with the drug efficacy on MTAP-deficient bladder cancers. *In silico* docking studies revealed that, unlike MTA, m-APTA interact inefficiently with adenosine A_1_ receptor, providing a plausible explanation of the superior safety profile of m-APTA.

**Conclusion:** m-APTA can significantly improve the translational potential of the NBA toxicity reduction strategy in selectively targeting MTAP-deficient cancers.

## Background

Nucleobase analogue (NBA) drugs, such as 5-florouracial (5-FU) and 6-thioguanine (6-TG), have been approved and used in the clinic since 1960s. These drugs are antimetabolites that directly interfere with DNA and/or RNA synthesis, thus inhibiting cell proliferation, causing cell death, and leading to tumor regression. Although NBA drugs are effective in tumor suppression, they are associated with a variety of adverse effects, such as severe anemia, leukopenia, thrombocytopenia, granulocytopenia, and gastrointestinal damage [1].

Methylthioadenosine phosphorylase (MTAP) is an enzyme that plays a major role in polyamine metabolism and is critical for the salvage of adenine and methionine [2]. MTAP catalyzes the hydrolytic cleavage of its only natural substrate *S*-methyl-5’-thioadenosine (MTA) to adenine and methylthioribose-1-phosphate. The *MTAP* gene, located on chromosome 9p21, is often co-deleted with *CDKN2A* with a frequency of up to 40% in certain human cancers, such as glioma and bladder cancers [3,4]. Deletion of the *MTAP* gene, as well as dysregulation of gene expression [5], leads to MTAP deficiency in human cancers. Although *MTAP* deficiency contributes to the tumor microenvironment that favors tumor progression, leading to poor disease outcome in multiple types of cancers [6–9], this functional loss of MTAP offers a unique opportunity for developing selective therapeutics. Various strategies have been pursued to exploit this unique opportunity, such as methionine deprivation therapy and *de novo* adenine synthesis inhibition. However, these strategies were later shown to be ineffective in clinical trials [10,11]. Synthetic lethality with MTAP-deficiency has also been proposed, taking advantage of the vulnerability of MTAP-deficient cells to protein arginine *N*-methyltransferase 5 inhibition [12,13]. A recent study reported that this strategy has shown a potential sign of clinical activity in patients [14].

In addition to the aforementioned strategies, co-administration of NBA drugs with MTA is an appealing approach for treating MTAP-deficient cancers, potentially affording specific toxicity reduction to patients. This strategy was first proposed by Lubin and Lubin in their seminal paper [15]. They have demonstrated *in vitro* that co-treatment of MTA and NBAs could protect MTAP-intact human fibroblasts from the cytotoxic effects of NBAs, whereas MTAP-deficient cancer cells were not protected. The protective effect is achieved by introducing MTA in excess, which is converted to adenine by MTAP. Consequently, the conversion of NBAs to toxic nucleotides is limited through depletion of their common substrate phosphoribosyl-5-pyrophosphate (PRPP), which is required for the conversion of nucleobase to nucleoside (**Fig. 1**). We and others have demonstrated that co-administration of 6-TG and MTA significantly inhibited MTAP-deficient cancers while the mouse hosts were well protected from 6-TG-induced toxicity [16,17]. Recently, Tang et al. demonstrated that co-administration of MTA with 2-fluoroadenine (2-FA), a highly toxic NBA, protected the mouse hosts from 2-FA-induced toxicity while 2-FA remained suppressive to MTAP-deficient tumors [18].

**Fig. 1.**
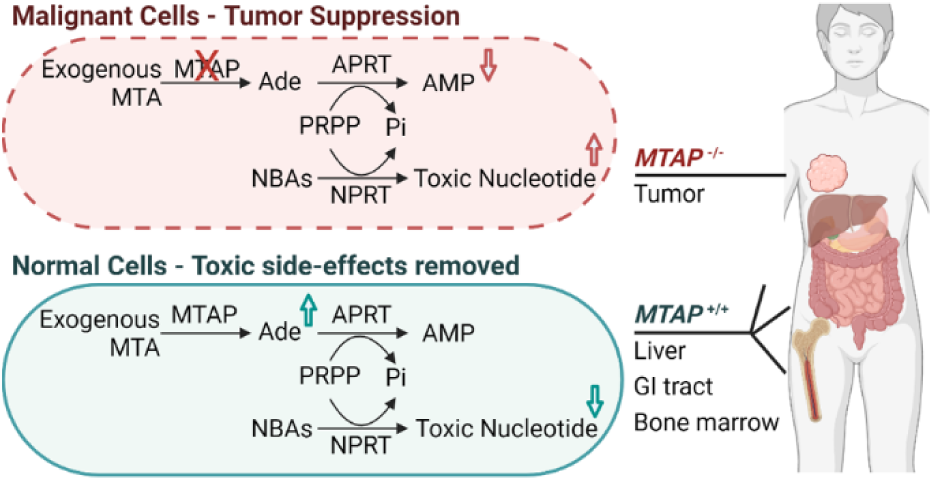
The treatment strategy and its mechanism of action. In normal cells, exogenously administrated MTA can be metabolized to adenine by the enzyme MTAP. This results in excessive adenine production that will deplete PRPP and prevent the conversion of NBAs to toxic nucleotides, thereby preventing their toxicity. In contrast, in MTAP-deficient malignant cells, this protective mechanism does not apply: no MTA is converted to adenine due to MTAP deficiency. Abbreviations: AMP, Adenosine monophosphate; APRT, adenine phosphoribosyltrasferase; MTA, 5’-deoxy-5’-methylthioadenosine; MTAP, MTA phosphorylase; NBA, nucleobase analogue; NPRT, nucleobase phosphoribosyltrasferase; Pi, inorganic phosphates; PRPP, phosphoribosyl pyrophosphate.

Among the clinically approved NBAs, 5-FU is commonly used to treat a wide range of cancers with well-studied mechanism of action and adverse effects [19]. Therefore, we set out to evaluate the utility of the MTA+5-FU combination in treating MTAP-deficient cancers *in vivo*. However, we noticed a dramatic decrease in body temperature (i.e. hypothermic effect) in mice following administration of MTA at the therapeutic dose, and this observation is consistent with a previous report [20]. Therefore, MTA is not readily translatable into clinical use. This undesirable effect of MTA is presumably due to the activation of the A_1_R in the brain and the heart [21]. We hypothesized that certain analogues of MTA could retain the protective effect against NBA toxicity while minimizing or eliminating the hypothermic effect. Here, we report the identification of an MTA analogue, m-APTA, that protects MTAP-intact cells when in use in combination with 5-FU without the induction of hypothermia. In addition, m-APTA does not reduce the efficacy of 5-FU. Furthermore, *In silico* docking experiments demonstrated that m-APTA had a favorable interaction with MTAP, but not with A_1_R.

## Materials and methods

### Chemicals

5-FU and 5’-deoxyadenosine (5’-dAdo) were obtained from Millipore-Sigma Canada Ltd. Adenosine was purchased from Chem-Impex (IL, United States), and MTA was bought from Abcam (Cambridge, UK). MTA analogues used for *in vitro* experiments were synthesized as described in the Supplementary Methods, while m-APTA used in animal studies was obtained from Toronto Research Chemicals Inc., Canada. Reagents used for HPLC studies were obtained from Fisher Scientific (NH, United States). For enzyme assays, recombinant human MTAP was purchased from Origene (MD, United States, catalogue no. TP306024), and microbial xanthine oxidase was obtained from Millipore-Sigma (MA, United States, catalogue no. X2252).

### Animal studies

Male NRG mice utilized in the experiments were bred in Animal Resource Centre, BC Cancer Research Institute, Vancouver, British Columbia, Canada. All experimental protocols were approved by the University of British Columbia Animal Care Committee (A19-0025).

Two bladder cancer PDX models, LTL-490 and LTL-543, obtained from the Living Tumor Laboratory (www.livingtumorlab.com) [22,23] were used for the efficacy studies. For detail experimental procedures, please refer to Supplementary Methods.

### Enzyme activity assays

Cleavage of MTA analogues by MTAP was assessed using a coupled enzyme assay adapted from protocols described by Kung *et al*. and Savarese *et al* [24,25]. Frozen stocks of MTAP and xanthine oxidase (XO) were thawed on ice. Working solutions were prepared in 1x PBS (pH 7.4, 0.1 M) and the microplate reader used for analysis was maintained at 37 °C. Test compounds (MTA, PTA, adenosine, and PTA derivatives **1**–**12**) were diluted from 100 mM DMSO stock solutions to provide working solutions of 0.5 mM.

The wells of a 96-well plate were partially filled with PBS buffer (50 µL/well), followed by XO working solution (10 µL of 0.05 U/µL XO). The substrate solution (20 µL of 0.5 mM test compound) was then added, followed by MTAP working solution (20 µL of 0.02 µg/µL MTAP). The final substrate concentration in each well was 0.1 mM. Control wells used for monitoring any background (non-enzymatic) hydrolysis contained additional PBS buffer in place of the MTAP working solution, such that the concentration of test compound and XO remained constant.

The appearance of adenine oxidation products (due to XO-promoted oxidation of released adenine) was monitored by absorbance at 305 nM, with readings collected at 5-minute intervals. The slope of the change in absorbance was used to assess the rate of cleavage; all data were normalized to the rate of MTA cleavage.

### Cell lines

HEK293 and 293T cells were cultured in DMEM medium (Hyclone, Washington, D.C., United States) supplemented with 10% FBS (GIBCO, Thermo Fisher Scientific, Waltham, MA). 253J cells were cultured in RPMI-1640 medium (GIBCO) supplemented with 10% FBS (GIBCO). Isogenic MTAP-intact cells, HEK:wt, and MTAP-deficient cells, HEK:ΔM, were generated from HEK293 cells. Please refer to Supplementary Methods for the detailed procedures of the generation and maintenance of these cell lines.

### *In vitro* efficacy assay

Cells were seeded on 96-well plates at a density of 5000 cells per well. Twenty-four hours after seeding, the medium was replaced with drug-containing medium. Different concentrations of 5-FU were prepared by serial dilution. Each dose was tested in triplicate for each experiment. Cells were cultured for 72 hours and then measured for cell growth using an MTS assay (CellTiter 96® AQ_ueous_ one solution cell proliferation assay, Promega). The IC_50_ values were generated from 3 independent experiments using Prism 8.0 (GraphPad, San Diego, CA). Statistical differences between the IC_50_ were analyzed by Prism 8.0 using one-way ANOVA followed by multiple comparison.

### *In silico* docking studies

The computer-added docking studies of the small molecules to the target protein were performed using the Glide module of the Schrodinger suite of programs [26]. The crystal structures of 5EUB of MTAP and 6D9H of A_1_R, respectively, were prepared and used for docking. The hydrophobicity of the small molecules was predicted using the Slog P algorism [27].

#### Data availability

This study includes no data deposited in external repositories.

## Results

### MTA alleviated 5-FU toxicity *in vivo* but induced adverse effects

We first investigated whether MTA could provide protection from 5-FU toxicity *in vivo*, as the protective effect on MTAP-intact cells was previously reported with *in vitro* studies [19]. NRG mice were treated with 5-FU at a dose of 120 mg/kg/week. After a three-dose treatment, 3 out of 4 mice treated with only 5-FU became moribund due to 5-FU toxicity, while all mice survived when MTA was co-administrated at 0.3 mmol/kg (dosage adapted from previous reports [16–18]) or at 0.2 mmol/kg (**Fig. 2A**). In addition, 5-FU-induced body weight loss was prevented by MTA at both doses (**Fig. 2B**). Diarrhea was observed in all mice treated with 5-FU alone, but not in the mice treated with 5-FU and MTA combined. These results confirmed that MTA was effective in reducing 5-FU toxicity *in vivo*. However, hypothermia was observed in mice administered with 0.2 or 0.3 mmol/kg of MTA (**Fig. 2C**). These mice exhibited substantial reduction in physical activity, and they were only responsive when physical stimuli were applied. Since the hypothermic effect of MTA is highly unfavorable for clinical use, we performed a titration to determine the MTA dose that would not induce hypothermia. After the administration of MTA at 0.2 or 0.3 mmol/kg, a drop of rectal temperature by 5.1±1.1 °C was observed while the hypothermic effect was absent at 0.1 mmol/kg (**Fig. 2C**). We then assessed whether MTA at 0.1 mmol/kg could protect animals from 5-FU toxicity. NRG mice were given the combination of 5-FU (80 mg/kg) and MTA weekly for 3 weeks. Blood was collected to evaluate the protective effect against the myelosuppressive effect of 5-FU. Our results showed that the protective effect of MTA on erythropoiesis was abolished when the dose of MTA was decreased to 0.1 mmol/kg while the toxicity was significantly alleviated at 0.3 mmol/kg (**Fig. 2D**). Taken together, our results showed that MTA could protect mouse hosts against 5-FU toxicity; however, the dosage of MTA required to achieve protection is associated with undesirable hypothermic effects. With a low MTA dose, the hypothermic effect could be prevented; however, the protective effect was lost at the lower dose. Thus, MTA is not a suitable protective agent for clinical use.

**Fig. 2.**
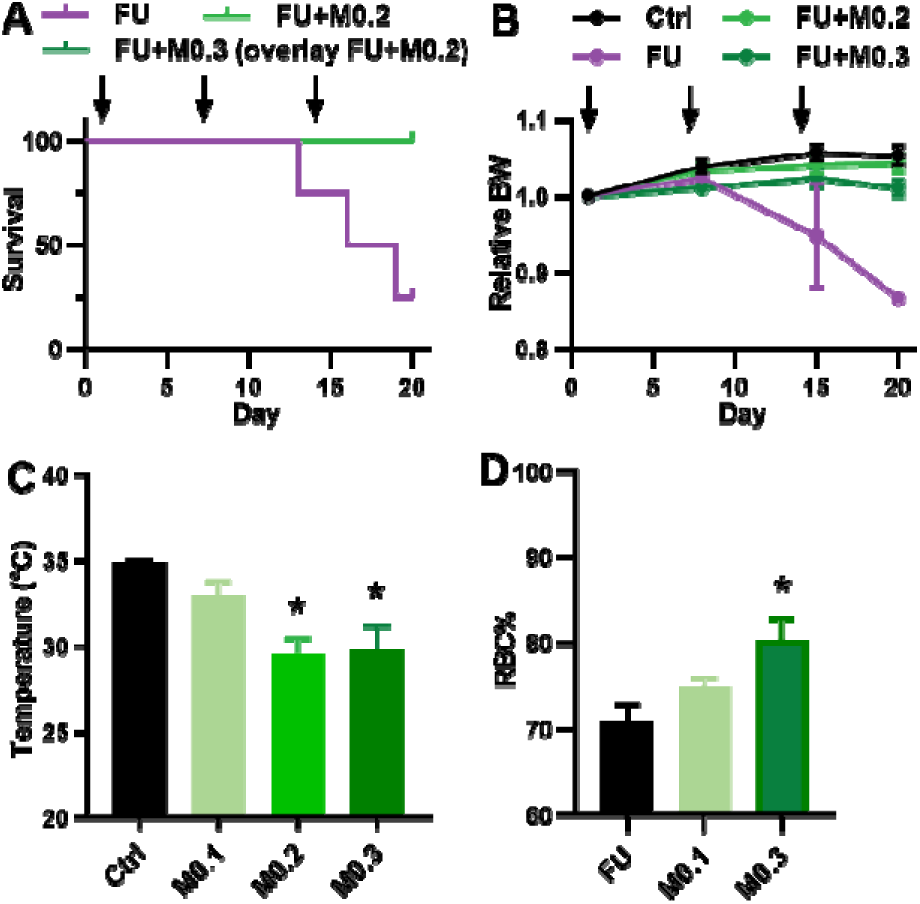
The effect of 5-FU +/- MTA on healthy mice. Male NRG mice (n=4) treated weekly with 5-FU at 120 mg/kg (FU) alone or in combination with MTA at 0.2 (M0.2) or 0.3 (M0.3) mmol/kg; control mice were treated with vehicle. Arrows indicate the day of treatment. (A) The survival curves of the mice in various groups are presented here. (B) Relative body weight was normalized to the original body weight of each mouse at day 0. Average relative body weights are presented with SEM. Note: body weight was averaged from 3 survived mice on Day 15, and 1 mouse on Day 20 for the 5-FU group. FU, 5-FU; M, MTA. (C) Average body temperature of male NRG mice (n=3) treated with MTA at different doses or the vehicle control (no protection). Rectal temperature was measured at 1 hour after the injections. Error bars indicate SEM.* *P*<0.05 compared with vehicle controls using one-way ANOVA followed by multiple comparisons analysis. (D) Male NRG mice (n=3) treated with 5-FU (80 mg/kg) alone or in combination with MTA at 0.1 (M0.1) or 0.3 (M0.3) mmol/kg weekly for 3 weeks. Four days after the 3^rd^ treatment, blood was collected for RBC counts. The RBC counts of each mouse were normalized to the original RBC count of each corresponding mouse within the treatment groups before the initiation of treatments. Error bars indicate the SEM. * *P*<0.05 compared to no rescue analyzed by one-way ANOVA followed by multiple comparisons analysis.

### Two phenylthioadenosine derivatives were identified as potential MTA substitutes

To translate this NBA toxicity reduction principle into the clinic, it is necessary to develop an alternative agent that functions similarly to MTA, yet without the adverse effects. We hypothesized that certain MTA analogues may exhibit these properties, and reasoned that the candidate should meet the following criteria: 1) it can be converted to adenine since this is essential for the protective mechanism; 2) it can protect MTAP-intact/normal tissues from 5-FU toxicity, 3) it does not bear the adverse effects of MTA at its effective dose, and 4) it does not interfere with the tumor suppression effect of 5-FU. To test the hypothesis, we started by identifying MTA analogues that were MTAP substrates with reasonable conversion rates for the production of adenine. A previous study demonstrated that substitution of the methyl group by a carboxylic aromatic ring (e.g. phenylthioadenosine, or PTA, **Fig. 3A**) retained reasonable conversion rate to adenine by MTAP [24]. A subset of 12 PTA derivatives (Fig. S1) were synthesized and evaluated, which were predicted to possess better water solubility for potential ease of downstream drug development, due to the inclusion of polar functional groups (refer to the Supplementary Methods for detailed synthetic protocols and characterization details). The ability of recombinant MTAP to cleave each of these molecules to adenine was assessed, since this is required for the protective mechanism (**Fig. 1**). The cleavage rate of each compound was normalized to that of MTA. At the same time, we confirmed that production of adenine in the absence of MTAP was negligible for each compound (<5%), indicating that none of the synthesized PTA derivatives were prone to non-enzymatic hydrolysis (**Fig. 3B**). Most significantly, two compounds, 5’-*S*-(4-aminophenyl)-5’-thioadenosine (m-APTA) and 5’-*S*-(4-aminophenyl)-5’-thioadenosine (p-APTA), exhibited improved adenine conversion rates relative to PTA. The relative rates of adenine production by MTAP from m-APTA and p-APTA were measured at 64% and 46% to that of MTA, respectively; in comparison, PTA had a relative conversion rate of 39% (**Fig. 3B**). Both m-APTA and p-APTA were therefore selected for further study.

**Fig. 3.**
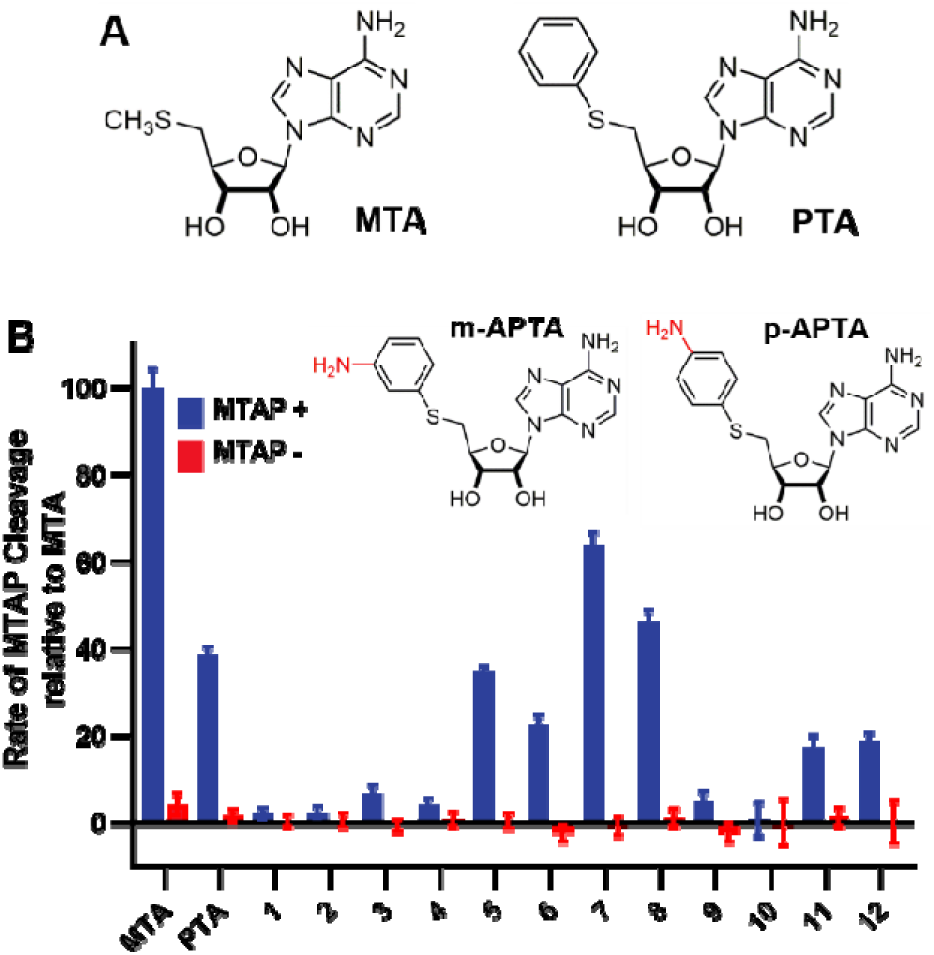
The structures and adenine conversion rates of MTA and its analogues. (A) Chemical structures of MTA and PTA. (B) Relative adenine conversion rates of selected MTA analogues at 0.1 mM in the presence (MTAP+) or absence (MTAP-) of MTAP. Error bars indicate SEM.

### m-APTA protected MTAP-intact but not MTAP-deficient cells from 5-FU cytotoxicity

To determine if the two candidates were functionally similar to MTA when in use in combination with 5-FU, we first subjected the agents to *in vitro* assessment. These candidates were found less toxic than MTA, for <5% of HEK293 cell proliferation was inhibited by m-APTA or p-APTA at up to 100 µM, whereas MTA resulted in a similar level of toxicity at 25 µM (Fig. S2). These concentrations were used for all subsequent studies to assess their protective effects.

Isogenic MTAP-intact (HEK:wt) and MTAP-deficient (HEK:ΔM) cell lines were generated (**Fig. 4A**) to investigate the efficacy and specificity of the protective effect of the candidates. Cells were treated with MTA, m-APTA, or p-APTA at the concentrations mentioned above in combination with 5-FU at various concentrations to generate dose-responsive curves for IC_50_ calculation. Our results showed that m-APTA protected HEK:wt cells from 5-FU-induced cytotoxicity by increasing the IC_50_ from 4.30±0.03 to 11.34±0.26 µM (**Fig. 4B, C**), although the protective effect was relatively smaller than that of MTA with IC_50_ of 23.15±0.84 µM (**Fig. 4C**). In contrast, p-APTA, the isomer of m-APTA, did not demonstrate significant protection, with IC_50_ of 5.99±0.20 µM (**Fig. 4B, C**). In addition, our results demonstrated that deletion of MTAP in HEK:ΔM cells abolished the protective effect of MTA or m-APTA against 5-FU-induced toxicity (**Fig. 4D**), confirming the specificity of the protective effect in MTAP-intact cells. We further confirmed the specificity of the protective effect of m-APTA by demonstrating that m-APTA could not protect bladder cancer cell lines with MTAP deficiency, including 253J (**Fig. 4E**) and RT112:wt (Fig. S3); and m-APTA was able to rescue bladder cancer cell lines with MTAP expression, including TCCSUP and HT1376 (Fig. S3). Taken together, our results demonstrated that, similar to MTA, m-APTA specifically protected MTAP-intact cells from 5-FU-induced cytotoxicity while leaving the MTAP-deficient cells unprotected. In contrast, p-APTA failed to demonstrate the protective effect, and was therefore excluded from subsequent *in vivo* investigation.

**Fig. 4.**
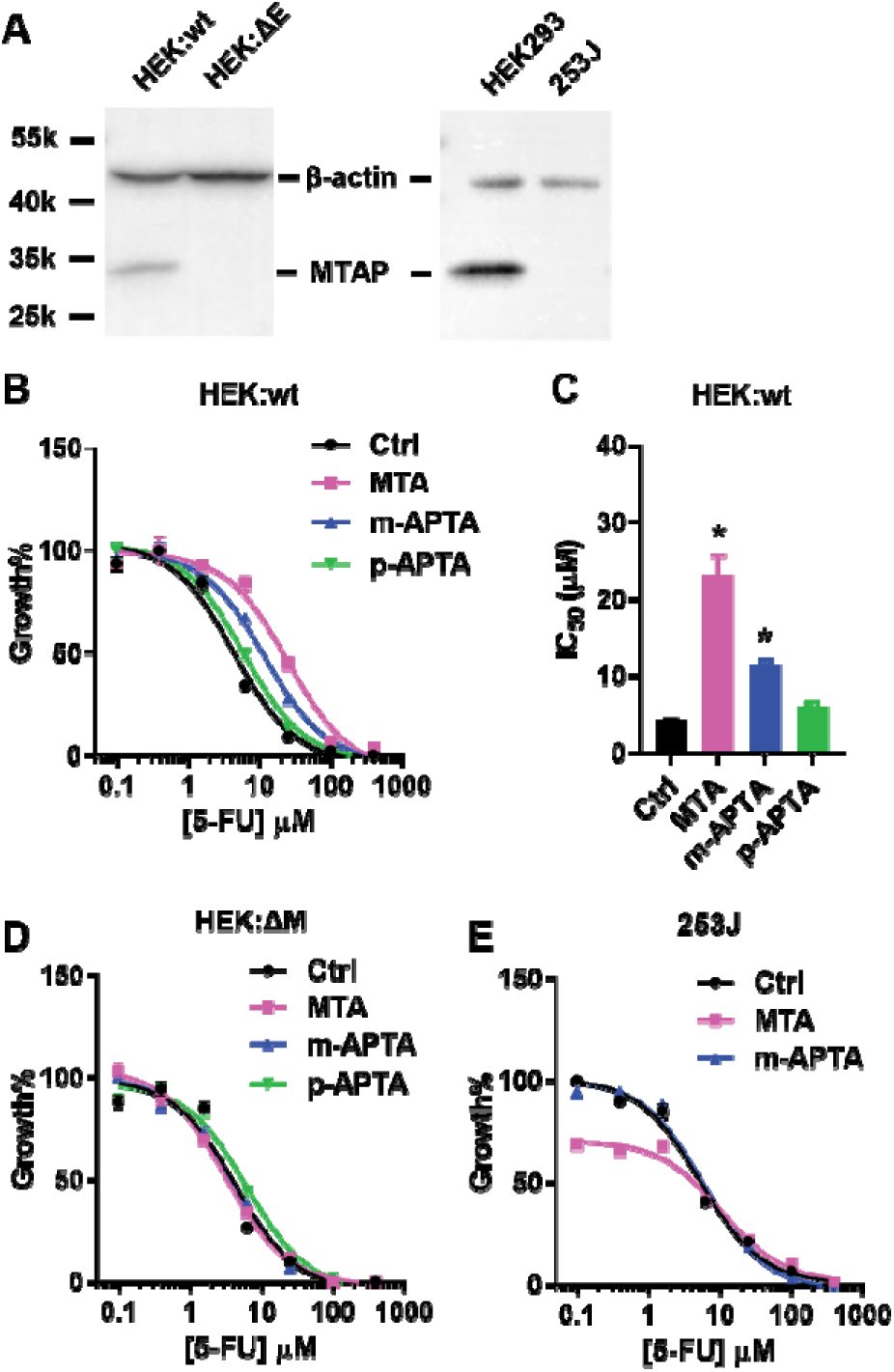
The effect of MTA analogues on 5-FU cytotoxicity. (A) Western blot with anti-MTAP of whole cell lysates from the indicated cell lines. HEK:wt and HEK:ΔM are isogenic cell lines with and without MTAP expression, respectively. 253J is a bladder cancer cell line that does not express MTAP. HEK293 cells are included as a control. (B) Dose-response curves of MTAP-positive HEK:wt cells to 5-FU alone (ctrl, black line) in addition to MTA (red), m-APTA (blue), or p-APTA (green). (C) IC_50_ values calculated from the dose-response curves from panel B. (D, E) Dose-response curves of MTAP-deficient HEK:ΔM (D) and 253J (E) to 5-FU (control) or 5-FU in combination with MTA, m-APTA, or p-APTA. Cells were treated by 5-FU at various doses (n=3). At 72 hours after treatment, cell viability was assessed by MTS assay. Error bars indicate SEM. * *P*<0.05 by one-way ANOVA followed by multiple comparisons analysis.

### m-APTA exhibited superior *in vivo* properties compared to MTA

We next evaluated the protective function of m-APTA *in vivo* and sought to determine whether it possessed the sedative effect associated with MTA. MTA and another MTA analogue 5’-dAdo were also included, since 5’-dAdo was previously proposed as an MTA substitute that demonstrated comparable protective effects against 5-FU-induced toxicity *in vitro* [15]. Healthy mice were given a combination treatment of 5-FU (80 mg/kg) and each protective agent (0.3 mmol/kg) once per week for 4 weeks. RBC counts were assessed as readouts of the protective function. During the course of the experiment, the RBC counts of the mice treated with 5-FU alone kept decreasing, and this was rescued by the presence of MTA, m-APTA, or 5’-dAdo (**Fig. 5A**). The final RBC counts of the 5-FU treated mice were decreased by ∼ 30%. The MTA or analogues supplemented groups rescued the RBC% by 10%, indicating that m-APTA equivalently protected erythropoiesis from 5-FU toxicity when compared to MTA and 5’-dAdo (**Fig. 5A**). Rectal temperature and behavior observations were recorded to evaluate sedative effects one hour following administration of the protective agents. Consistently, mice treated with MTA exhibited decreased rectal temperature by 5.8±0.2°C versus the control mice (**Fig. 5B**). The mice exhibited lack of activity but responded to physical stimuli. Mice subjected to 5’-dAdo suffered more severe rectal temperature decrease by 7.9±0.1°C versus the control (**Fig. 5B**). These mice lost the responsiveness to physical stimuli. Thus, 5’-dAdo is not a good candidate to replace MTA. In contrast, the protective dose of m-APTA did not trigger hypothermia (**Fig. 5B**). Taken together, m-APTA maintained the protective ability of MTA without the adverse effects when given at a therapeutic dose, fulfilling the first two criteria of the candidate to replace MTA.

**Fig. 5.**
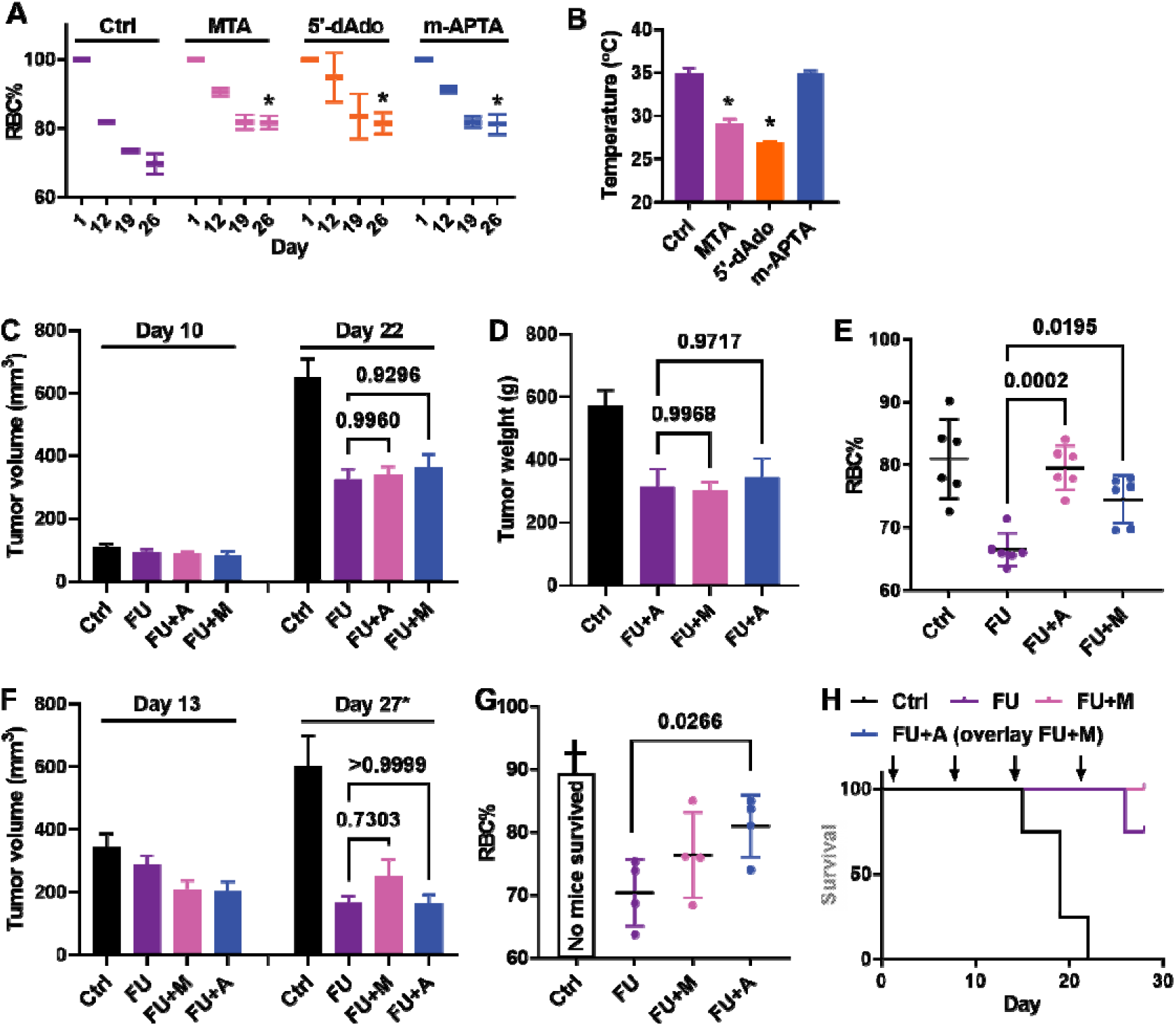
The host protection and tumor suppression efficacy of the combination treatment of 5-FU and m-APTA. (A, B) Male NRG mice (n=3) treated with 5-FU alone (Ctrl) or in combination with MTA, m-APTA, or 5’-dAdo weekly for 4 weeks. (A) Four days after the 2^nd^, 3^rd^, and 4^th^ treatment, blood was collected for RBC counts. RBC counts% is normalized with the RBC counts before treatment. Error bars indicate the SEM. * *P*<0.05 compared to control on Day 26 by one-way ANOVA followed by multiple comparisons analysis. (B) Rectal temperature was measured at 1 hour after the injections. The averages of the measurements are presented here. Error bars are SD. * *P*<0.05 compared to control by one-way ANOVA followed by multiple comparisons analysis. (C-E) Male NRG mice bearing LTL-543 PDXs were randomized into 4 groups (n=6) and subjected to 5-FU treatment alone (FU), in the combination with MTA (FU+M) or m-APTA (FU+A), or with the vehicle control (Ctrl, sham treatment). Animals were treated weekly for 3 weeks and euthanized on Day 22. The average tumor volumes (C) and tumor weights (D) as well as the RBC counts% (E) are presented. RBC counts% is normalized to the RBC counts before treatment. Error bars indicate SD. Statistics were analyzed by one-way ANOVA followed by multiple comparison analysis. (F-H) Male NRG mice bearing LTL-490 PDXs were randomized into 4 groups (n=4) and subjected to 5-FU treatment alone (FU), or in the combination with MTA (FU+M) or m-APTA (FU+A) in comparison to the vehicle group (Ctrl). Animals were treated weekly for 4 weeks and euthanized on Day 28 or at the humane endpoint. Average tumor volumes (F), RBC counts% (G), and survival (H) are presented here. (F) * Individual tumor volumes in the control group were measured on Day 17, 19, 21 when mice were euthanized at the humane endpoint, while the tumor volumes of the other groups were measured on day 27. (G) RBC counts% is normalized with the RBC counts before treatment. Statistics were analyzed by one-way ANOVA followed by multiple comparison analysis. (H) All mice survived in groups FU+M and FU+A.

### m-APTA did not interfere with 5-FU anti-tumour activity in MTAP-deficient bladder cancer PDX models

For m-APTA to be used in combination with 5-FU, it cannot interfere with the tumor suppressive effect of 5-FU on MTAP-deficient cancer. We selected bladder cancers as our initial study focus, since 5-FU is a treatment option for bladder cancers and MTAP loss is frequently observed (∼1 in 4 cases, Fig. S4) [28–31]. The effect of 5-FU + m-APTA was evaluated in two bladder cancer PDX models, LTL-490 and LTL-543, developed in the Living Tumor Lab [22]. Both PDX models were derived from patients with muscle-invasive bladder cancers harboring *MTAP* deletion, which was confirmed by whole genome DNA sequencing (Supplementary Methods, Fig. S5). The original tumor sample of LTL-543 was derived from treatment-naïve tumor sample, while LTL-490 was subjected to neoadjuvant chemotherapy. Detailed information of the PDX models is summarized in **Table 1**. The small pieces of PDX seeds were implanted under the renal capsule of NRG mice. When the tumor size reached about 70 mm^3^ for LTL-543 or 100 mm^3^ for LTL-490, the mice were randomized into four groups to receive the following treatments once per week for 3 weeks for LTL543 or 4 weeks for LTL-490: i) vehicle control, ii) 5-FU alone, iii) 5-FU + MTA and iv) 5-FU + m-APTA. RBC counts were measured to assess the protective effect. After 5-FU treatment, the average tumor size was significantly reduced by 50 ± 9% in volume on day 22 for LTL-543 (**Fig. 5C**) and 63 ± 14% in volume on day 27 for LTL-490 (**Fig. 5F**). Moreover, the final average tumor weight was significantly reduced by 45% ± 12% for LTL-543 (**Fig. 5D**). As expected, treatment of 5-FU alone significantly reduced RBC counts. Consistent with the observations on the healthy mice, supplementing m-APTA significantly rescued the reduction of RBC counts caused by 5-FU (**Fig. 5E, G**). Importantly, neither MTA nor m-APTA compromised 5-FU efficacy: the tumor weight and volume was comparable between 5-FU alone group and the combination groups along the treatment course (**Fig. 5C, D, F**). In addition, LTL-490-bearing mice treated with the vehicle control did not survive beyond three weeks after implant while the mice treated with 5-FU alone or in combination with m-APTA survived much longer (**Fig. 5H**). Therefore, m-APTA is able to exert its protective effect on erythropoiesis without compromising the efficacy of 5-FU on the MTAP-deficient PDXs.

**Table 1.** The genetic background and growth characteristics of the bladder PDX (*40*).

| PDX model | DNA_seq | Patient | Pathologic diagnosis | Pathologic stage/grading | Neoadjuvant chemotherapy |
| --- | --- | --- | --- | --- | --- |
| LTL-490 | *MTAP*<sup>-/-</sup> | 67/male | Urothelial Carcinoma | pT4bN0Mx/G3 | Gemcitabine/Cisplatin |
| LTL-543 | *MTAP*<sup>-/-</sup> | 70/male | Urothelial Carcinoma | pT4aN3Mx/G3 | Naïve |

### In silico modeling showed an unfavorable interaction between m-APTA and A_1_R

To gain further insight into the *in vivo* behavior of m-APTA, we compared the ability of MTA and m-APTA to bind to MTAP and A_1_R using a proven computer-aided drug design (CADD) platform [32]. We first compared the interaction of the APTA isomers with MTAP. We found that the docking position of m-APTA in MTAP was generally the same as that of MTA, with the adenine side embedded in the active site of the enzyme (**Fig. 6A, B**). In contrast, the p-APTA docks into MTAP in an opposite direction (not shown), with the adenine branch directing away from the active site of MTAP. It is plausible that lower adenine conversion rate compared to m-APTA was due to the difference in binding orientation relative to MTA.

**Fig. 6.**
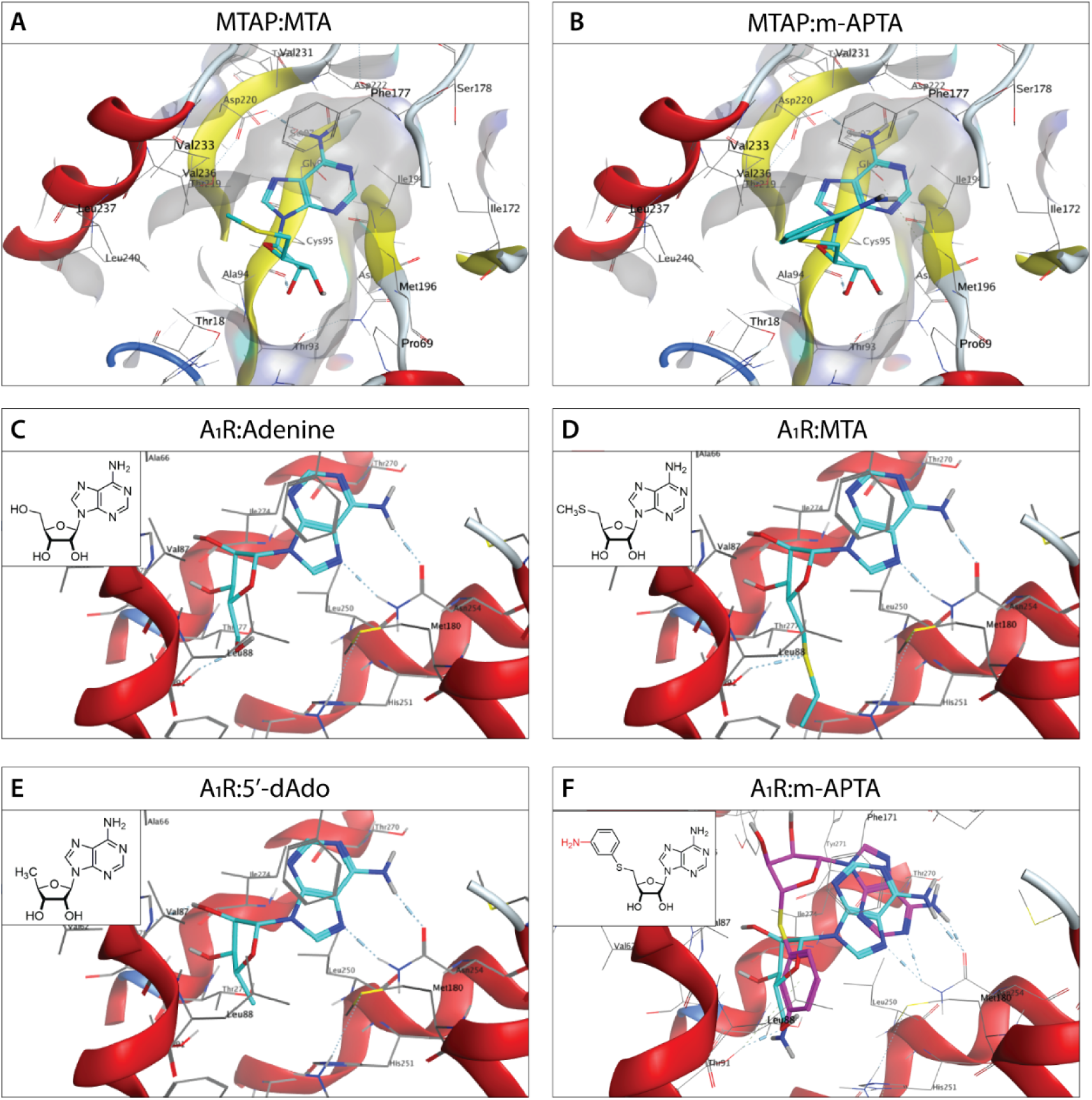
Computer-generated molecular docking of small molecules to MTAP or adenosine A_1_ receptor (A_1_R). Molecular docking of MTAP (reference PDB:5EUB) with MTA (A) and m-APTA (B). Molecular docking of A_1_R (reference PDB: 6D9H) with adenosine (C and cyan in F), MTA (D), 5’dAdo (E), and m-APTA (pink) (F)

Our docking analysis further showed that MTA and 5’-dAdo interact with A_1_R in the same configuration as that observed for adenosine, the natural agonist of A_1_R (**Fig. 6C-E**). This presumably results in the hypothermic effect observed with these agents [33]. In contrast, molecular docking revealed that substitution of the 5’-methyl group in MTA by a functionalized aromatic ring meant that m-APTA could no longer interact with A_1_R (**Fig. 6F**). The binding selectivity of A_1_R with MTA, 5’-dAdo, or m-APTA are therefore consistent with the loss of sedative effect of m-APTA.

## Discussion

NBAs are effective in killing cancer cells but do not spare normal proliferating cells. In principle, the utility of NBAs can be improved through methods that widen the therapeutic windows of these drugs, including the reduction of NBA toxicity to normal tissues or the enhancement of their cytotoxic effects on the cancer cells. Particularly for 5-FU, combination approaches with leucovorin, thymidine, or uridine have been assessed [34–37]. However, only leucovorin was shown to be effective in the clinical setting. In addition to these strategies, the co-administration of MTA and NBAs for the treatment of MTAP-deficient cancer was proposed [15]. The efficacy and specificity of this strategy has been well-documented by various groups with *in vitro* and *in vivo* studies [15–18]. However, we have discovered and are now reporting here for the first time that the effective level of MTA induces unwanted suppression of body temperature and sedation, which can engender significant health risks to cancer patients. To address this limitation, we report the discovery of m-APTA, the first-in-field solution to enable the safe application of this combination therapy, thus significantly enhancing its translational value.

MTA is the natural substrate of MTAP and, therefore, has a relatively high conversion rate in the MTAP-mediated production of adenine. However, this property does not necessarily make MTA the best drug candidate. In our study, it is interesting to note that MTA and m-APTA demonstrated equivalent protective effects *in vivo* at the same dose (Fig. 5) while MTA exhibited a superior in vitro protection (Fig. 4C). This may be attributable to the slower, and perhaps optimal, conversion rate of m-APTA to adenine (Fig 3). Given the same levels of *in vivo* protection produced by m-APTA and MTA when administered at the same dose, we speculate that although exogenously administered MTA can exert its protective effect due to its efficient conversion to adenine, both MTA and adenine are eliminated from the system quickly after administration [38–40]. In contrast, the slower adenine conversion rate of m-APTA may allow for longer *in vivo* exposure to adenine at a lower level, thus prolonging the protective window, and eventually contributing to an equivalent level of protection. These results also suggested that simply pursuing a high adenine conversion rate may not lead to the ideal *in vivo* protection; instead, a moderate conversion rate (with prolonged conversion) is probably sufficient to produce enough adenine. It is also important to note that exposure to adenine can be indirectly prolonged by modified pharmacokinetics of the MTA analogue, and this can be achieved by chemical modification or of the use of various drug delivery systems. Therefore, the considerations of both the conversion rate and the pharmacokinetics of the candidate may further improve the *in vivo* protective effect of m-APTA.

The utility of this toxicity reduction strategy hinges on the presence of MTAP in the normal cells while the tumors are MTAP-deficient. Genomic loss of *MTAP* is a passenger event due to its genomic proximity to *CDKN2A* [41,42], which encodes the tumor suppressor p16^INK4a^ that is often lost in various cancers. As for bladder cancers, loss of chromosome 9p21, which carries *CDKN2A* and *MTAP*, is an early event [43–46]. Therefore, there is a high probability that selected cancer cell clones during tumor progression will carry this genetic lesion. This, in turn, leads to a high probability that the pattern of *MTAP* loss will be more homogeneous in bladder cancers. In line with this, lack of heterogeneity of MTAP expression in MTAP-deficient cancers has been demonstrated in non-small cell lung carcinoma [47] and nasopharyngeal carcinoma [48], as also noted by Tang et al [18]. Such homogeneous loss of MTAP enhances the likelihood of success of the combination therapy.

With the identification of m-APTA, we posit that this toxicity reduction strategy can afford to treat patients with MTAP-deficient using more treatment cycles and/or higher doses. This may potentially improve overall treatment efficacy. Although the utility of m-APTA was demonstrated with 5-FU in this study, the application scope of m-APTA is not limited to combination with NBAs used under existing treatment protocols. Because the toxicity reduction mechanism involves the specific production of adenine through MTAP, which is applicable to other NBAs, we hypothesize that m-APTA can be combined with a variety of novel NBAs, such as 2,6-diaminopurine and 2-FA [15,49]. In addition, the m-APTA + NBA combination may be extended to treat cancer types that are not currently indicated. In some cancers, such as prostate cancer, no NBA has been approved for clinical use due to lack of efficacy at the tolerated dosage [50]. However, our previous studies have demonstrated that in combination with MTA, 6-TG can suppress MTAP-deficient castration-resistant prostate cancers (*MTAP* deletions occur in about 25% of prostate cancer cases) [17]. Discovery of m-APTA, therefore, allows the exploration of the full potential of NBAs.

## Conclusions

In this report, we have identified m-APTA as a superior substitute for MTA, for co-administration with NBAs. This compound fulfilled each of our pre-determined criterion: 1) it can be converted to adenine through the action of MTAP, 2) it functions similarly to MTA, protecting MTAP-intact cells against NBA-induced toxicity, 3) its administration at therapeutically efficacious concentrations did not induce any of the adverse effects seen with MTA, and 4) it does not interfere with the tumor suppression effect of the NBA drug (i.e. 5-FU). Because m-APTA does not induce sedative effects at its effective dosage, it has considerable merit as a safer substitution for MTA to be administered to cancer patients. The favorable properties of m-APTA have significantly increased the translational potential of the combination treatment strategy with NBAs. In addition, our findings strongly support additional studies aimed at using this combination in treating cancers with MTAP deficiency.

## List of abbreviations

2-FA: 2-fluoroadenine
5-FU: 5-florouracial
5’-dAdo: 5’-deoxyadenosine
6-TG: 6-thioguanine (6-TG)
A_1_R: Adenosine A_1_ receptor
i.p.: Intraperitoneal
m-APTA: 5’-*S*-(4-aminophenyl)-5’-thioadenosine
MTA: *S*-methyl-5’-thioadenosine
MTAP: Methylthioadenosine phosphorylase
NBA: Nucleobase analogue
p-APTA: 5’-*S*-(4-aminophenyl)-5’-thioadenosine
PDX: Patient-derived xenograft
PRPP: Phosphoribosyl-5-pyrophosphate
PTA: Phenylthioadenosine
RBC: Red blood cell
XO: Xanthine oxidase

## Declarations

### Ethics approval and consent to participate

Consented patient tumor samples received for the development of the PDX models were covered by human ethics protocols H04-60131, H09-01628, and H18-01520 approved by the Research Ethics Board at BC Cancer and University of British Columbia.

### Consent for publication

Not applicable

### Availability of data and materials

All data generated or analysed during this study are included in this published article and its supplementary information files.

### Author contributions

SZ and YW designed the study, with support from NKYW. SZ directed the study and performed the majority of the experiments. HX developed the PDX models and performed part of the efficacy studies. JD performed the cytotoxicity studies. TD, IB and JW synthesized the PTA derivatives and performed the enzyme assay. FB and AC performed the molecular docking analyses. SA, SV, YL, and CC performed the DNA-sequencing data analyses. SZ, NKYW, and YW wrote the manuscript.

## Acknowledgements

We would like to thank all members of the Living Tumor Laboratory for technical support and helpful discussions. The authors wish to acknowledge Canada’s Michael Smith Genome Sciences Centre, Vancouver, Canada for the whole genome sequencing of PDX models.

---

**Fig. S1.**
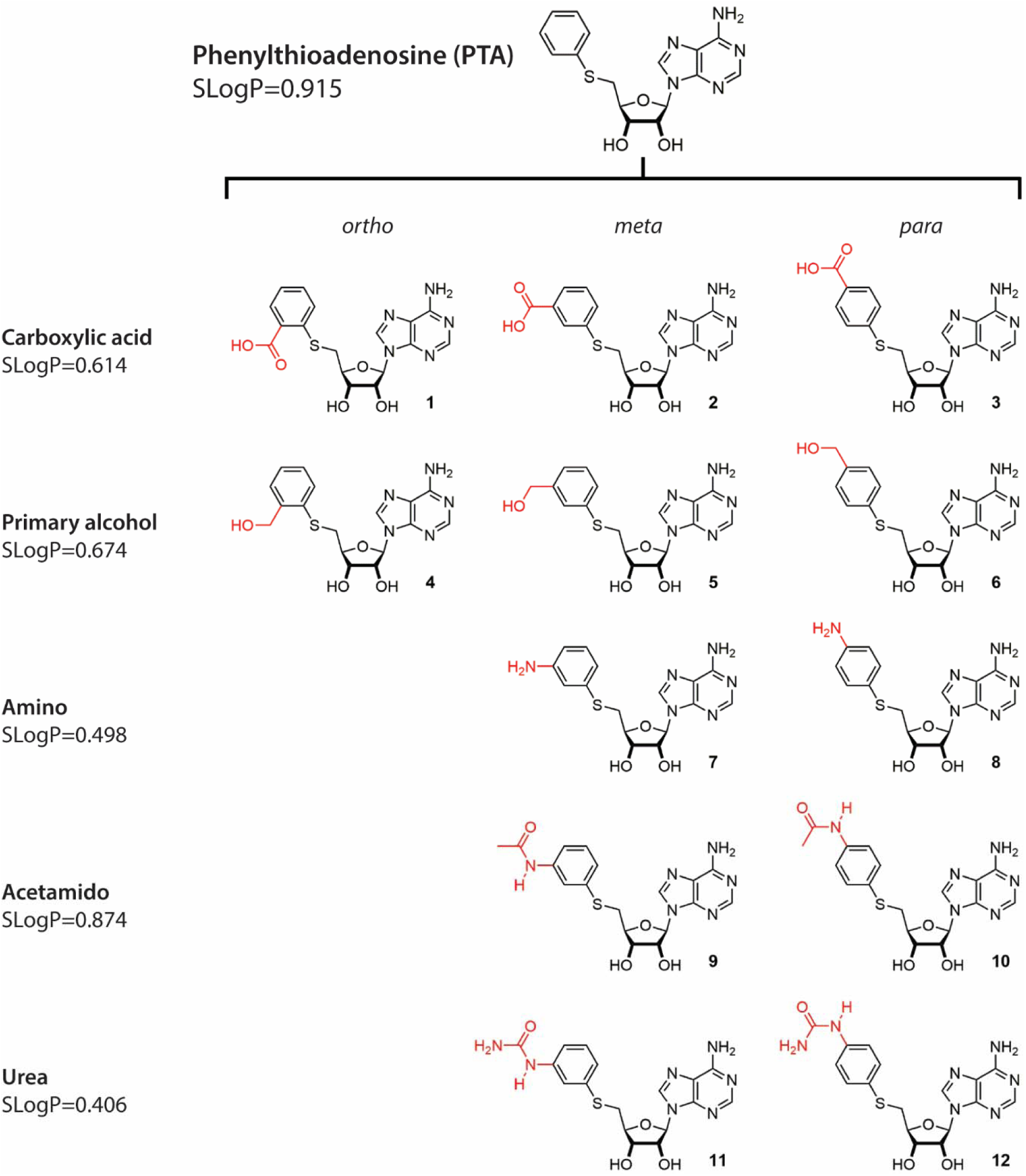
Phenylthioadenosine derivatives. Phenylthioadenosine derivatives containing polar functional groups were generated to increase water solubility. Because *ortho*-alcohol **4** was a poorer MTAP substrate than *meta*- and *para*-alcohols **5** and **6**, *ortho*-amino derivatives were not pursued. Water solubility is indicated by SLogP.

**Fig. S2.**
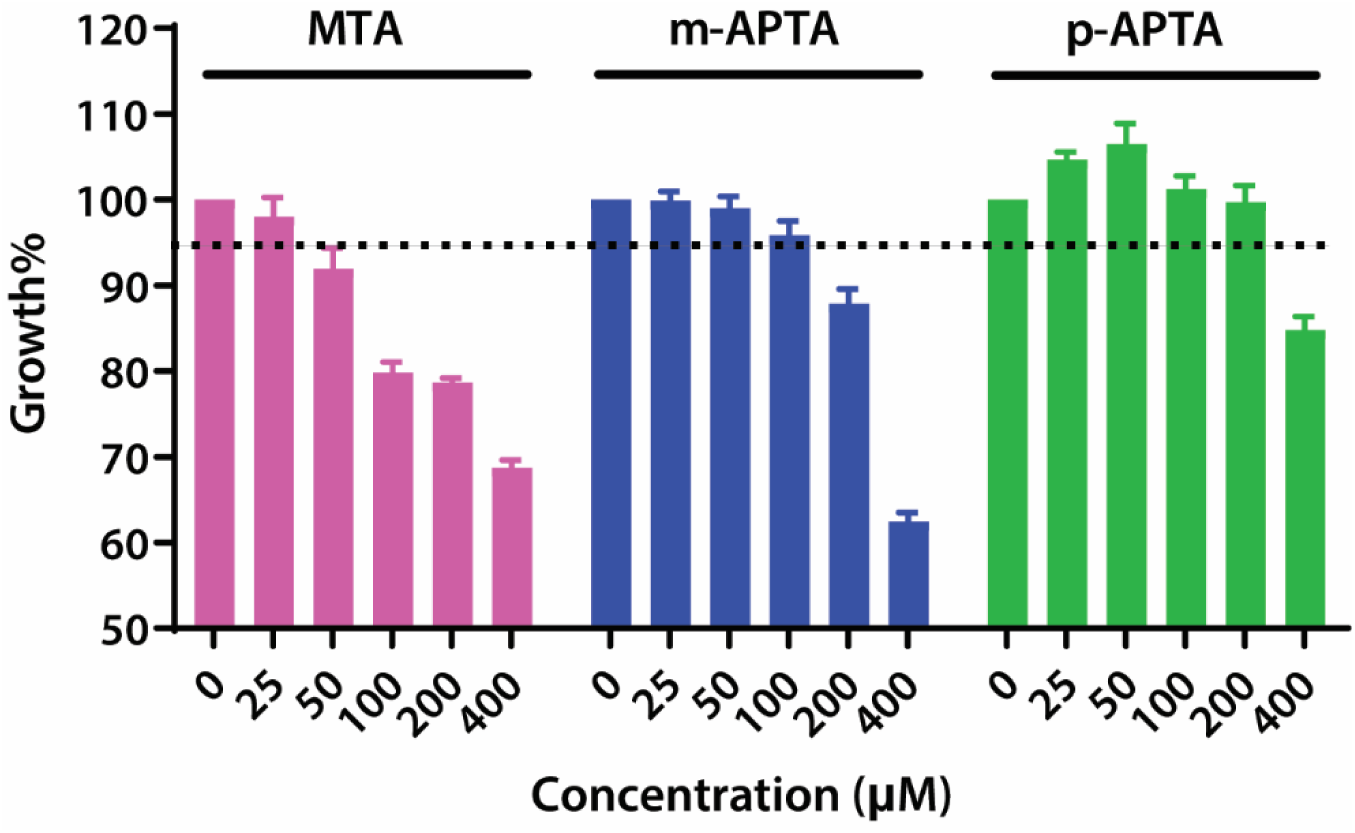
Cell growth inhibition induced by MTA or its analogues on HEK293. Cells were treated by MTA, m-APTA, or p-APTA at various concentrations (n=3). At 72 hours after treatment, cell viability was assessed by MTS assay. The cell growth% represents the relative cell growth to the vehicle control. Error bars indicate SEM.

**Fig. S3.**
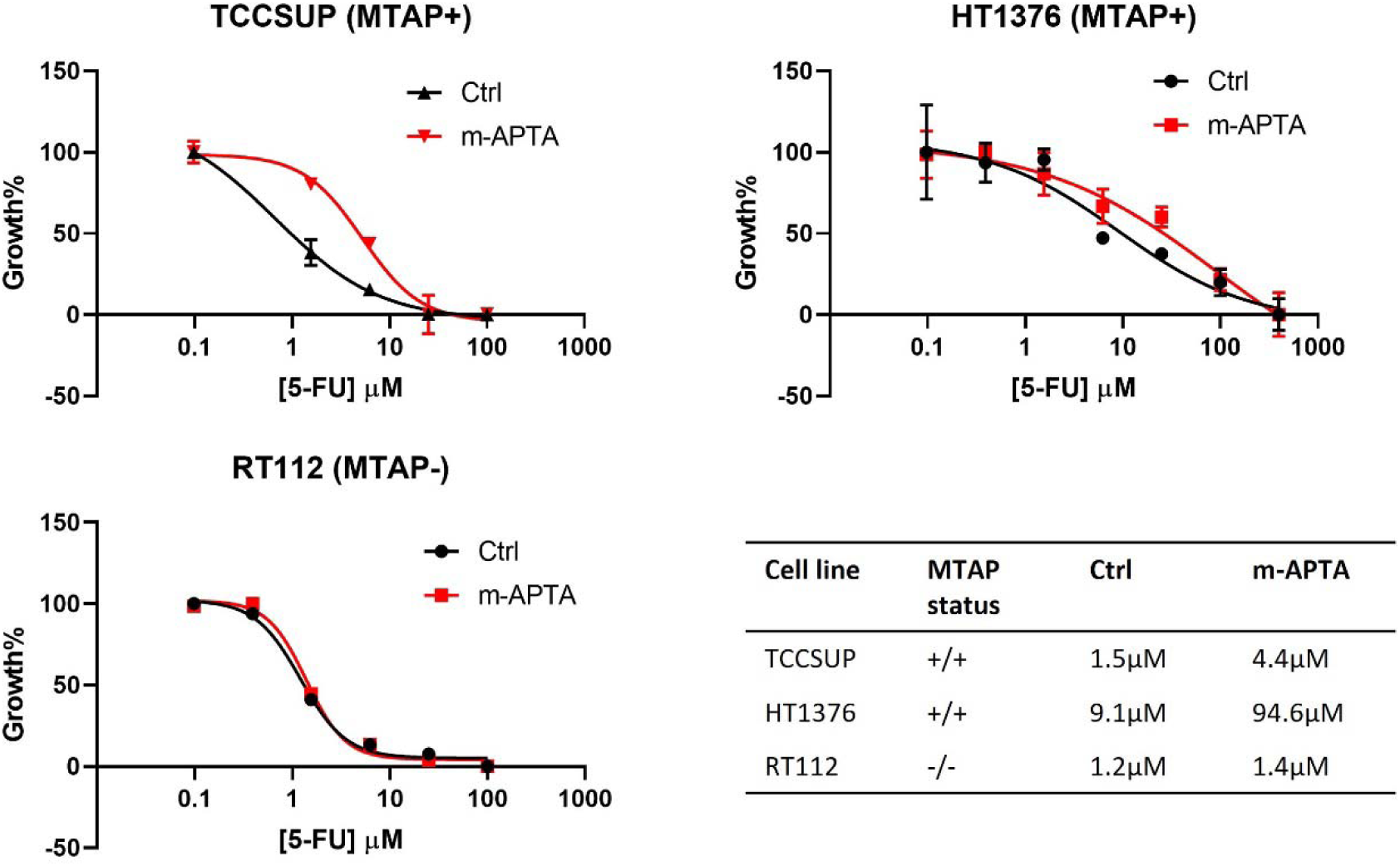
The effect of m-APTA on 5-FU cytotoxicity in bladder cancer cell lines. HT1376, TCCSUP, and RT112 cell lines were cultured in MEM medium (GIBCO) supplemented with sodium pyruvate (GBICO), non-essential amino acids (GIBCO), and 10% FBS (GIBCO). Cells were treated by 5-FU at various doses with or without the combination of m-APTA at 100 µM (n=3). At 72 hours after treatment, cell viability was assessed by MTS assay. IC_50_ values were calculated from the dose-response curves for the individual cell lines.

**Fig. 4.**
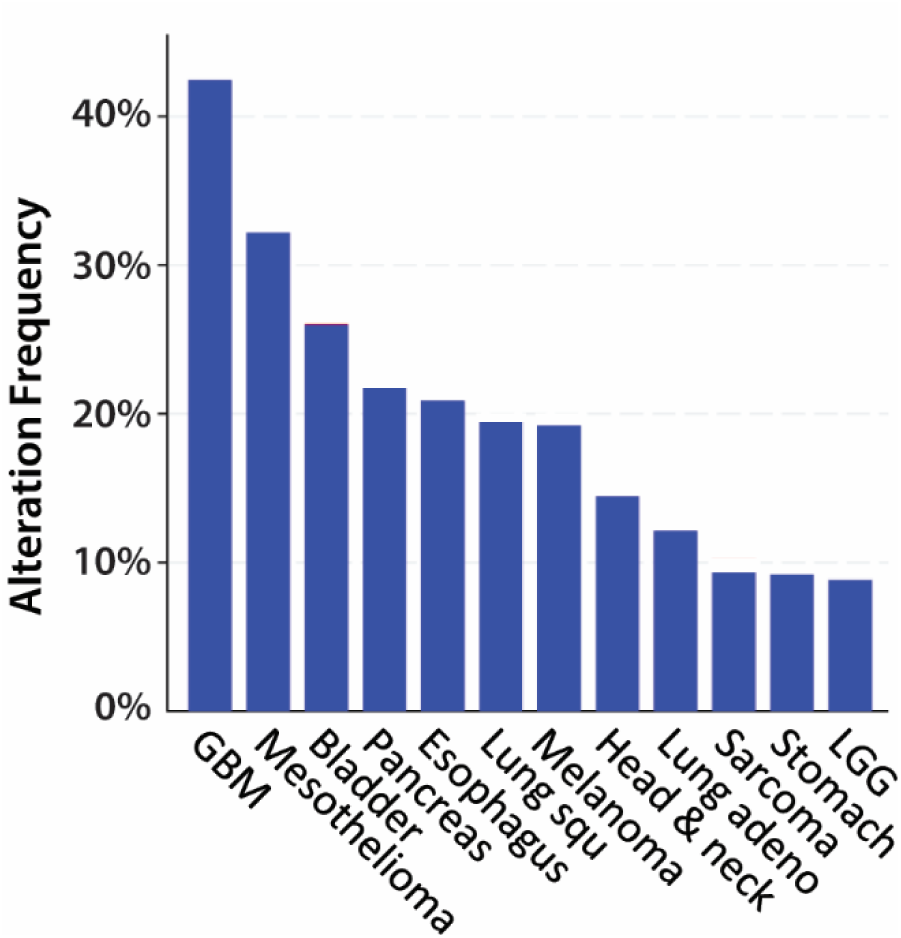
Genetic statuses of *MTAP* in different cancers. The frequencies of MTAP homozygous deletion in a variety of malignancies were obtained from cBioPortal. Thirty-two TCGA PanCancer Atlas Studies of different cancer types were included. This collection comprised 10953 patients, among which 1013 patient tumors had copy number alteration of *MTAP* – 978 were homozygous deletion. The cancer types reported with > 5% of homozygous deletion of MTAP are displayed. GBM, glioblastoma; LGG, low-grade glioma.

**Fig. S5.**
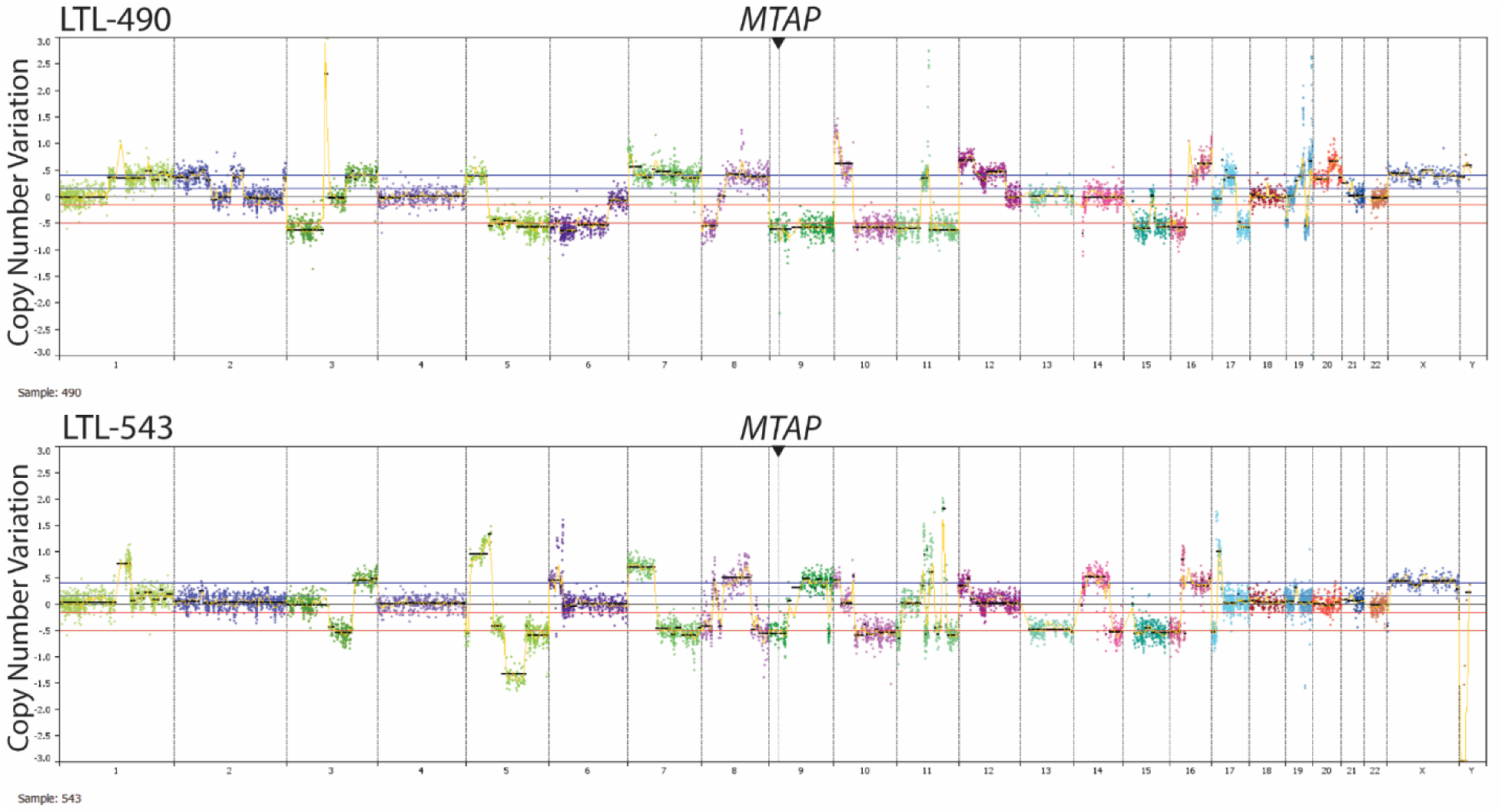
*MTAP* deletion in bladder PDX models. Copy number variations of LTL-490 and LTL-543 tumour genomes were generated by whole genome sequencing. Triangles indicate the position of MTAP on the chromosome 9.

## Supplementary File 1. Supplementary Methods

### Recovery of patient-derived xenograft tumor seed and conduction of mouse studies

Tumor seeds kept under liquid nitrogen were recovered from storage. Pieces of tissues were grafted under the renal capsules of NRG mice for the recovery and expansion of cancer tissue. After 2 months, the PDX tissues were harvested, cut into small pieces (1×2×3 mm^3^), and re-grafted into the subrenal capsule site of NRG mice for the efficacy studies. The tumor volumes at the middle time point during the treatment course were measure by ultrasound (Vevo 3100, FUJIFILM VisualSonics, Inc.), and the volumes were generated by the software Vevo LAB 2.1.0 (FUJIFILM VisualSonics, Inc.). Since the tumor cells of the PDXs still maintain typical bladder epithelial features and developed fluid-filled cysts, the tumor volumes were corrected by subtracting the volumes of the cavities when measured by ultrasound. For LTL-543, tumor volume could not be corrected for the cavity volume at the last time point because the volume was measured by a caliper. Therefore, the tumor weight after depleting the encapsulated fluid was also measured to quantify the tumor growth at the last time point. The tumor volumes at the end of the experiment were measured by caliper after excision, and the tumor volume was calculated using the formula L×W×H×0.52. The maximal permitted tumor size was 1500 mm^3^ by the ethics committee, and all tumor sizes were below the limit.

The combination treatment was administered to bladder cancer PDX-bearing mice. Each of the protective agents, namely MTA, 5’-dAdo, or m-APTA, was first dissolved in DMSO and then diluted in PBS before intraperitoneal (i.p.) injection. One hour after the injection, animals were subjected to i.p. injection of 5-FU dissolved in PBS. Mice were treated by the combination once per week for 3 or 4 weeks, as indicated. Rectal temperature was measured 1 hour after injection of the protective agents. The physical activity of the mice was monitored 1 hour after the mice were dosed. For mice lacking physical activity, the responsiveness of the mice was assessed following physical stimuli to the areas of head, both flanks, and tail of the mice. Body weight and general health was monitored daily. In some mice, red blood cell (RBC) counts were measured to assess the effect of 5-FU on erythropoiesis. After treatment, blood was collected 4 days after each 5-FU treatment, which is within the window of death and re-population of the proliferating cells in the bone marrow (28). Tail vein blood was collected into EDTA-coated tubes, kept on ice, and subjected to blood counting within an hour using Vet ABC haematology analyzer (scil animal care company GmbH, Barrie, ON). For each mouse, RBC count from the blood collected one day before the first treatment was set as the baseline to assess the treatment-induced changes of RBC counts. The mice were euthanized at the end of the experiment or when the humane endpoint was reached, including body weight loss >20% and tumor volume >5% of the body weight. One-way ANOVA followed by multiple comparison or *t* test was performed analyzed the differences among groups using Prism 8.0 (GraphPad, San Diego, CA).

### Generation and maintenance of isogenic MTAP-intact and MTAP-deficient cells

The HEK:ΔM cell line was generated through biallelic deletion of MTAP from HEK293 cells using CRISPR-Cas9 technology (31,32). The single guidance sequence targeting exon 5 of MTAP (CACCGGAAGGACTGAGGTCTCATAG) was annealed with the reverse complementary sequence (AAACCTATGAGACCTCAGTCCTTCC), and they were inserted into the lentiCRISPRv2 plasmid. The newly generated plasmid was co-transfected with psPAX2 and pVSVg into HEK293T cells for viral packaging, and the virus-containing supernatent was then used to infect HEK293 cells. Infected cells were diluted to 0.5 cells per 100µl and cultured in puromycin-supplemented medium on 96-well plates for single colony screening. The isolated colonies were expanded, and the gene deletion was validated by Western Blot using MTAP antibody (11475-1-AP, Proteintech). In the in vitro efficacy studies, HEK:ΔM (MTAP-deficient) was compared with HEK:wt (MTAP-intact) cells. HEK:wt cell line was generated by the same procedure except that empty lenti-CRISPRv2 plasmid was cotransfected with viral packaging plasmids. HEK:wt and HEK:ΔM cell lines were then maintained in DMEM medium (Hyclone) supplemented with 10% FBS (GIBCO) and 0.4 µg/ml puromycin (GIBCO).

### Synthesis of phenylthioadenosine derivatives

m-APTA (**7**) and p-APTA (**8**) were synthesized by addition of 3-aminothiphenol and 4-aminothiphenol, respectively, to 5’-chloro-5’-deoxyadenosine, which in turn was prepared through treatment of adenosine with thionyl chloride as described previously (24,25). Amides **9**– **10** were prepared similarly but using the corresponding acetamidothiophenol as the nucleophile. Analogues **1**–**6** were synthesized using a related approach, except that 5’-tosyladenosine was used as the electrophile, and the carboxylic acid and alcohol groups were accessed from the corresponding ester by saponification or reduction. Ureas **11**–**12** were prepared by derivatization of **7** and **8** with potassium cyanate. Synthetic and characterization details for all compounds are included in the Additional file 1.

### Synthesis of m-APTA and p-APTA from 5*’*-chloro-5*’*-deoxyadenosine

*5’-chloro-5’-deoxyadenosine.* Under an argon atmosphere, adenosine (15 mmol) was stirred at 0 °C in a mixture of acetonitrile (60 mL) and pyridine (32 mmol) for 10 minutes. Thionyl chloride (74 mmol) was added dropwise, and the mixture was stirred for 4 hours at 0 °C, followed by a further 16 hours at room temperature. The resulting yellow emulsion was filtered and washed with acetonitrile (2 x 20 mL). The precipitate was dissolved in methanol (60 mL) and aqueous ammonium hydroxide (2 M, 40 mL), and the resulting solution was stirred for 1 hour at room temperature, before being concentrated *in vacuo*. The crude product was dissolved in water (80 mL) at 70 °C. The solution was cooled to room temperature and then left to sit at 4 °C for 16 h to encourage crystallization. The crystals were filtered, washed with cold water, and dried under vacuum to provide the desired product as a white crystalline solid (3.98 g; 93% yield). ^1^H NMR (300 MHz, DMSO-d_6_): δ 8.34 (s, 1H), 8.16 (s, 1H), 7.31 (br s, 2H), 5.93 (d, *J* = 5.7 Hz, 1H), 5.61 (d, *J* = 6.9 Hz, 1H), 5.47 (d, *J* = 5.3 Hz, 1H), 4.75 (m, 1H), 4.23 (m, 1H), 4.09 (m, 1H), 3.95 (dd, *J* = 11.5, 5.02 Hz, 1H), 3.84 (dd, *J* = 11.5, 6.4 Hz, 1H); ^13^C NMR (75 MHz, DMSO-d_6_): 156.1, 152.8, 149.5, 139.8, 119.2, 87.5, 83.7, 72.7, 71.3, 44.8.

5*’*-*S*-(3-aminophenyl)-5*’*-thioadenosine *(m-APTA; **7**).* 3-Aminothiophenol (3.8 mmol) was dissolved in dry DMF (10 mL) and placed under an argon atmosphere. Potassium *tert*-butoxide (1M in THF; 4.1 mmol) was added, and the mixture was stirred at room temperature for 30 minutes, after which a solution of 5*’*-chloro-5*’*-deoxyadenosine (1.9 mmol) in DMF (2 mL) was quantitatively injected. Stirring was continued at 40 °C for 16 hours, after which the mixture was neutralized with 10% aqueous HCl, and concentrated *in vacuo*. The crude product was recrystallized from 1:1 methanol:water (10 mL) by dissolving the material at 70 °C and then cooling the resulting solution at 4 °C for 16 hours. The precipitate was filtered, washed with cold water and dichloromethane (2 x 20 mL) to afford the desired product as a tan solid (497 mg, 70% yield). ^1^H (500 MHz; DMSO-d_6_): δ 8.35 (s, 1H), 8.16 (s, 1H), 7.30 (br s, 2H), 6.93 (m, 1H), 6.55 (m, 1H), 6.47 (m, 1H), 6.38 (m, 1H), 5.89 (d, *J* = 5.7 Hz, 1H), 5.48 (br, 2H), 5.31 (br s, 2H), 4.81 (m, 1H), 4.17 (m, 1H), 4.00 (m, 1H), 3.32 (dd, *J* = 14.1, 6.4 Hz, 1H), 3.19 (dd, *J* = 13.8, 7.0 Hz, 1H); ^13^C (125 MHz DMSO-d_6_): δ 156.0 (C), 152.6 (CH), 149.5 (C), 149.2 (C), 139.9 (CH), 135.7 (C), 129.4 (CH), 119.2 (C), 115.3 (CH), 113.2 (CH), 111.7 (CH), 87.3 (CH), 82.9 (CH), 72.6 (CH), 72.5 (CH), 35.1 (CH_2_); IR (neat, cm^-1^) 3280 (m), 3146 (m), 3132 (m), 2918 (m), 1680 (s), 1652 (s), 1596 (s), 1578 (s), 1480 (s), 1423 (m), 1342 (m), 1297 (s), 1208 (m), 1129 (m), 1089 (s), 1040 (s), 990 (m), 832 (m), 745 (s), 713 (s), 685 (s), 639 (s), 593 (s), 542 (s); HRMS (ESI) calculated for [C_16_H_18_N_6_O_3_S + H^+^] 375.1234, found 375.1230; M.P. 85–88 °C.

5*’*-*S*-(4-aminophenyl)-5*’*-thioadenosine *(p-APTA; 8).* 4-Aminothiophenol (7.6 mmol) was dissolved in dry DMF (10 mL) and placed under an argon atmosphere. Potassium *tert*-butoxide (1M in THF; 8.3 mmol) was added, and mixture was stirred at room temperature for 30 minutes, after which a solution of 5*’*-chloro-5*’*-deoxyadenosine (2.5 mmol) in DMF (2 mL) was quantitatively injected. Stirring was continued at 40 °C for 16 hours, after which the mixture was neutralized with 10% aqueous HCl, and concentrated *in vacuo*. The crude product was recrystallized from 1:1 methanol:water (10 mL) by dissolving the material at 70 °C and then cooling the resulting solution at 4 °C for 16 hours. The precipitate was filtered, washed with cold water and dichloromethane (2 x 20 mL) to afford the desired product as a tan solid (856 mg, 91% yield). ^1^ H (500 MHz DMSO-d_6_): δ 8.35 (s, 1H), 8.15 (s, 1H), 7.30 (br s, 2H), 7.11 (m, 2H), 6.51 (m, 2H), 5.87 (d, *J* = 5.7 Hz, 1H), 5.52 (br s, 1H), 5.36 (br s, 1H), 4.79 (m, 1H), 4.13 (m, 1H), 3.90 (m, 1H), 3.12 (dd, *J* = 13.6, 6.5 Hz, 1H), 2.99 (dd, *J* = 13.6, 6.9 Hz, 1H); ^13^C (125 MHz DMSO-d_6_): δ 156.0 (C), 152.6 (CH), 149.5 (C), 148.5 (C), 139.9 (CH), 134.3 (CH), 133.7 (CH), 119.2 (C), 118.3 (C), 114.4 (CH), 114.2 (CH), 87.1 (CH), 83.0 (CH), 72.5 (CH), 72.5 (C), 38.8 (CH_2_); IR (neat, cm^-1^) 3276 (m), 3132 (m), 2897 (m), 1674 (m), 1651 (s), 1600 (s), 1575 (s), 1496 (s), 1421 (m), 1341 (m), 1294 (s), 1127 (s), 1098 (s), 1036 (s), 817 (s), 717 (s), 630 (s), 513 (s); HRMS (ESI) calculated for [C_16_H_18_N_6_O_3_S + H^+^] 375.1234, found 375.1231; M.P. 96–99°C.

### Synthesis of PTA derivatives 1–6 from 5’-*O*-tosyladenosine

*5’-S-[2-(methoxycarbonyl)phenyl]-5’-thioadenosine.* Under an argon atmosphere, methyl 2-mercaptobenzoate (1.50 mmol) was dissolved in DMF (10 mL). NaH (1.94 mmol) was added, and the mixture was stirred for 30 minutes, becoming bright yellow. 5’-*O*-Tosyladenosine (1.25 mmol) was added to the stirring solution. The resulting mixture was stirred for 16 hours, affording a brown solution. The product was extracted into ethyl acetate (50 mL), washed 3 times with distilled water (15 mL), and concentrated *in vacuo*. The residue was purified by recrystallization: the product was dissolved in methanol (10 mL) at 60 °C, cooled to room temperature, then incubated at 4 °C for 16 hours to allow crystals to form. These were filtered, washed with cold methanol followed by cold dichloromethane, and then dried under vacuum to afford the desired product as a white powder in 84% yield. ^1^H (500 MHz; DMSO-d_6_): δ 8.36 (s, 1H), 8.15 (s, 1H), 7.85 (d, *J* = 7.9 Hz, 1H), 7.51 (t, *J* = 9.2 Hz, 1H), 7.50 (d, *J* = 6.9 Hz, 1H), 7.29 (br s, 2H), 7.24 (td, *J* = 6.86, 2.2 Hz, 1H), 5.91 (d, *J* = 5.7 Hz, 1H), 5.54 (d, *J* = 5.9 Hz, 1H), 5.40 (d, *J* = 4.9 Hz, 1H), 4.82 (m, 1H), 4.23 (m, 1H), 4.07 (m, 1H), 3.82 (s, 3H), 3.43 (dd, *J* = 13.9, 5.5 Hz, 1H), 3.34 (dd, *J* = 14.1, 7.3 Hz, 1H); ^13^C (125 MHz; DMSO-d_6_): δ 166.1 (C), 156.1 (C), 152.6 (CH), 149.5 (C), 139.9 (CH), 139.9 (C), 132.6 (CH), 130.7 (CH), 127.6 (C), 126.1 (CH), 124.3 (CH), 119.2 (C), 87.5 (CH), 82.4 (CH), 73.0 (CH), 72.6 (CH), 52.1 (CH_3_), 34.2 (CH_2_); IR (neat, cm^-1^) 3339 (w), 3172 (m), 2930 (w), 1706 (s), 1671 (s), 1611 (m), 1429 (m), 1303 (m), 1246 (s), 1230 (s), 1115 (m), 1021 (s), 787 (s), 720 (s), 706 (s), 688 (s), 660 (s); HRMS (ESI) calculated for [C_18_H_19_N_5_O_5_S + H^+^] 418.1180, found 418.1180; M.P. 117–119 °C.

*5’-S-[3-(methoxycarbonyl)phenyl]-5’-thioadenosine.* Under an argon atmosphere, methyl 3-mercaptobenzoate (1.50 mmol) was dissolved in DMF (10 mL). NaH (1.94 mmol) was added, and the mixture was stirred for 30 minutes, becoming bright yellow. 5’-*O*-Tosyladenosine (1.25 mmol) was added to the stirring solution. The resulting mixture was stirred for 16 hours, affording a brown solution. The product was extracted into ethyl acetate (50 mL), washed 3 times with distilled water (15 mL), and concentrated *in vacuo*. The residue was purified by recrystallization: the product was dissolved in methanol (10 mL) at 60 °C, cooled to room temperature, then incubated at 4 °C for 16 hours to allow crystals to form. These were filtered, washed with cold methanol followed by cold dichloromethane, and then dried under vacuum to afford the desired product as a white powder in 46% yield. ^1^H (500 MHz; DMSO-d_6_): δ 8.32 (s, 1H), 8.14 (s, 1H), 7.87 (t, *J* = 1.7 Hz, 1H), 7.74 (dt, *J* = 7.8, 1.4 Hz, 1H), 7.63 (m, 1H), 7.43 (t, *J* = 7.9 Hz, 1H), 7.28 (br s, 2H), 5.89 (d, *J* = 5.9 Hz, 1H), 5.52 (d, *J* = 6.2 Hz, 1H), 5.38 (d, *J* = 4.9 Hz, 1H), 4.82 (m, 1H), 4.21 (m, 1H), 4.02 (m, 1H), 3.82 (s, 3H), 3.49 (dd, *J* = 13.6, 5.6 Hz, 1H), 3.39 (dd, *J* = 13.6, 7.2 Hz, 1H); ^13^C (125 MHz; DMSO): δ 165.7(C), 156.1 (C), 152.6 (CH), 149.4 (C), 139.9 (CH), 137.0 (C), 132.7 (CH), 130.4 (C), 129.4 (CH), 128.2 (CH), 126.4 (CH), 119.2 (C), 87.5 (CH), 82.8 (CH), 72.7 (CH), 72.5 (CH), 52.3 (CH_3_), 35.0 (CH_2_); IR (neat, cm^-1^) 3328 (m), 3184 (m), 2925 (m), 1717 (m), 1640 (s), 1599 (m), 1574 (m), 1418 (m), 1287 (s), 1262 (m), 1123 (s), 1029 (s), 747 (s), 646 (s); HRMS (ESI) calculated for [C_18_H_19_N_5_O_5_S + H^+^] 418.1180, found 418.1180; M.P. 94–97 °C.

*5’-S-[4-(methoxycarbonyl)phenyl]-5’-thioadenosine.* Under an argon atmosphere, methyl 4-mercaptobenzoate (1.50 mmol) was dissolved in DMF (10 mL). NaH (1.94 mmol) was added, and the mixture was stirred for 30 minutes, becoming bright yellow. 5’-*O*-Tosyladenosine (1.25 mmol) was added to the stirring solution. The resulting mixture was stirred for 16 hours, affording a brown solution. The product was extracted into ethyl acetate (50 mL), washed 3 times with distilled water (15 mL), and concentrated *in vacuo*. The residue was purified by recrystallization: the product was dissolved in methanol (10 mL) at 60 °C, cooled to room temperature, then incubated at 4 °C for 16 hours to allow crystals to form. These were filtered, washed with cold methanol followed by cold dichloromethane, and then dried under vacuum to afford the desired product as a white powder in 69% yield. ^1^H (500 MHz; DMSO-d_6_): δ 8.34 (s, 1H), 8.15 (s, 1H), 7.82 (d, *J* = 8.7 Hz, 2H), 7.43 (d, *J* = 8.4 Hz, 2H), 7.28 (br s, 2H), 5.90 (d, *J* = 5.8 Hz, 1H), 5.54 (d, *J* = 6.0 Hz, 1H), 5.40 (d, *J* = 4.9 Hz, 1H), 4.82 (m, 1H), 4.23 (m, 1H), 4.06 (m, 1H), 3.82 (s, 3H), 3.53 (dd, *J* = 13.8, 5.5 Hz, 1H), 3.43 (dd, *J* = 13.8, 7.4 Hz, 1H); ^13^C (125 MHz; DMSO): 165.8 (C), 156.1 (C), 152.7 (CH), 149.3 (C), 143.4 (C), 139.9 (CH), 129.5(CH), 126.3 (CH), 126.1 (C), 119.2 (C), 87.5 (CH), 82.6 (CH), 72.7 (CH), 72.6 (CH), 52.0 (CH_3_), 33.8 (CH_2_); IR (neat, cm^-1^) 3389 (m), 3327 (m), 3114 (m), 2946 (w), 1708 (m), 1686 (m), 1663 (s), 1596 (s), 1571 (m), 1333 (m), 1290 (s), 1119 (s), 1092 (s), 1013 (s), 760 (s); HRMS (ESI) calculated for [C_18_H_19_N_5_O_5_S + H^+^] 418.1180, found 418.1180; M.P. 210–211 °C.

*5’-S-(2-carboxyphenyl)-5’-thioadenosine (**1**).* 5’-*S*-[2-(methoxycarbonyl)phenyl]-5’-thioadenosine (0.5 mmol) was stirred in NaOH (10 wt%, 3 mL) and water (3 mL) for 24 hours. The mixture was neutralized with 10% aqueous HCl, concentrated *in vacuo*, and dried to form a white solid. The crude carboxylic acid was triturated with a small amount of water to remove residual salt, prior to enzymatic analysis. ^1^H (500 MHz; DMSO-d_6_): δ 8.65 (s, 1H), 8.43 (s, 1H), 7.86 (m, 1H), 7.47 (m, 2H), 7.20 (m, 1H), 5.96 (d, *J* = 5.7 Hz, 1H), 4.76 (m, 1H), 4.23 (m, 1H), 4.10 (m, 1H), 3.43 (dd, *J* = 13.6, 5.6 Hz, 1H), 3.29 (dd, *J* = 13.5, 7.5 Hz, 1H); ^13^C (125 MHz; DMSO-d_6_): δ 167.3 (C), 148.7 (C), 141.6 (CH), 140.0 (C), 132.2 (CH), 130.9 (CH), 128.3 (C), 128.0 (CH), 125.6 (CH), 125.5 (CH), 124.0 (CH), 118.8 (C), 87.6 (CH), 82.7 (CH), 73.1 (CH), 72.9 (CH), 33.9 (CH_2_); IR (neat, cm^-1^) 3318 (m), 3147 (m), 2903 (m), 1700 (s), 1416 (m), 1310 (m), 1254 (s), 1084 (s), 1044 (s), 950 (m), 831 (m), 748 (s), 719 (s), 637 (m), 536 (m); HRMS (ESI) calculated for [C_17_H_17_N_5_O_5_S + H^+^] 404.1023, found 404.1023; M.P. 150–153 °C.

*5’-S-(3-carboxyphenyl)-5’-thioadenosine (**2**).* 5’-*S*-[3-(methoxycarbonyl)phenyl]-5’-thioadenosine (0.5 mmol) was stirred in NaOH (10 wt%, 3 mL) and water (3 mL) for 24 hours. The mixture was neutralized with 10% aqueous HCl, concentrated *in vacuo*, and dried to form a white solid. The crude carboxylic acid was triturated with a small amount of water to remove residual salt, prior to enzymatic analysis. ^1^H (500 MHz; DMSO-d_6_): δ 8.36 (s, 1H), 8.15 (s, 1H), 7.87 (m, 1H), 7.69 (m, 1H), 7.31 (m, 1H), 7.30 (br s, 2H), 7.22 (m, 1H), 5.89 (d, *J* = 6.0 Hz, 1H), 4.80 (m, 1H), 4.20 (m, 1H), 4.01 (m, 1H), 3.41 (dd, *J* = 13.9, 6.0 Hz, 1H), 3.28 (dd, *J* = 13.7, 7.2 Hz, 1H); ^13^C (125 MHz; DMSO-d_6_): δ 169.5 (C), 156.0 (C), 152.6 (CH), 149.4 (C), 140.8 (C), 139.9 (CH), 134.3 (C), 129.2 (CH), 128.7 (CH), 128.0 (CH), 126.7 (CH), 125.5 (C), 119.1 (C), 87.3 (CH), 82.7 (CH), 72.7 (CH), 72.6 (CH), 35.5 (CH_2_); IR (neat, cm^-1^) 3304 (w), 3179 (m), 1644 (m), 1595 (s), 1550 (s), 1477 (m), 1379 (s), 1044 (m), 763 (s), 646 (s); HRMS (ESI) calculated for [C_17_H_17_N_5_O_5_S + H^+^] 404.1023, found 404.1023; M.P. 210–215 °C.

*5’-S-(4-carboxyphenyl)-5’-thioadenosine (**3**).* 5’-*S*-[4-(methoxycarbonyl)phenyl]-5’-thioadenosine (0.5 mmol) was stirred in NaOH (10 wt%, 3 mL) and water (3 mL) for 24 hours. The mixture was neutralized with 10% aqueous HCl, concentrated *in vacuo*, and dried to form a white solid. The crude carboxylic acid was triturated with a small amount of water to remove residual salt, prior to enzymatic analysis. ^1^H (500 MHz; DMSO-d_6_): δ 8.36 (s, 1H), 8.15 (s, 1H), 7.80 (d, *J* = 8.5 Hz, 2H), 7.37 (d, *J* = 8.5 Hz, 2H), 7.30 (br s, 2H), 5.90 (d, *J* = 5.8 Hz, 1H), 4.81 (m, 1H), 4.22 (m, 1H), 4.05 (m, 1H), 3.53 (dd, *J* = 13.8, 5.6 Hz, 1H), 3.39 (dd, *J* = 13.8, 7.6 Hz, 1H); ^13^C (125 MHz; DMSO-d_6_): δ 167.4 (C), 156.0 (C), 152.6 (CH), 149.4 (C), 141.6 (C), 139.9 (CH), 129.7(CH), 126.3 (CH), 119.1 (C), 87.5 (CH), 82.6 (CH), 72.7 (CH), 72.6 (CH), 34.1 (CH_2_); IR (neat, cm^-1^) 3325 (m), 2968 (w), 1704 (s), 1590 (m), 1488 (m), 1391 (s), 1308 (m), 1183 (s), 1085 (s), 1052 (s), 1009 (m), 950 (m), 772 (s), 725 (s), 627 (s), 515 (s); HRMS (ESI) calculated for [C_17_H_17_N_5_O_5_S + H^+^] 404.1023, found 404.1023; M.P. 193–195 °C.

*5’-S-[2-(hydroxymethyl)phenyl]-5’-thioadenosine (**4**).* Under an argon atmosphere, 5’-*S*-[2-(methoxycarbonyl)phenyl]-5’-thioadenosine (0.4 mmol) was stirred in dioxane (4 mL), forming an emulsion. LiAlH_4_ (2.4 mmol) was added, and the mixture was stirred at 50 °C for 3 hours.

The reaction mixture was neutralized by the addition of ethyl acetate (4 mL) and HCl (1 M, 2 mL), and then extracted into water (2 mL) three times. The grey solid was filtered out of the aqueous layer, washed with water, and dried *in vacuo* to afford the desired product as a clear, colorless oil in 69% yield. ^1^H (500 MHz; DMSO-d_6_): δ 8.35 (s, 1H), 8.14 (m, 1H), 7.44 (m, 1H), 7.39 (m, 1H), 7.27 (m, 2H), 7.20 (m, 2H), 5.89 (m, 1H), 5.59 (br s, 1H), 5.47 (br s, 1H), 5.30 (br s, 1H), 4.80 (m, 1H), 4.53 (s, 2H), 4.20 (m, 1H), 4.00 (m, 1H). ^13^C (125 MHz; DMSO-d_6_): δ 155.9 (C), 152.6 (CH), 149.4 (C), 141.3 (C), 139.9 (CH), 132.9 (C), 128.1 (CH), 127.2 (CH), 126.7 (CH), 125.7 (CH), 119.0 (C), 87.3 (CH), 82.8 (CH), 72.7 (CH), 72.6 (CH), 60.5 (CH_2_), 35.4 (CH_2_); IR (neat, cm^-1^) 3345 (s), 2964 (w), 2924 (w), 2854 (w), 2257 (w), 1646 (s), 1600 (m), 1475 (m), 1420 (m), 1337 (m), 1237 (m), 1090 (m), 1047 (s), 1025 (s), 988 (s), 828 (s), 766 (s), 504 (s); HRMS (ESI) calculated for [C_17_H_19_N_5_O_4_S + H^+^] 390.1231, found 390.1232.

*5’-S-[3-(hydroxymethyl)phenyl]-5’-thioadenosine (**5**).* Under an argon atmosphere, 5’-*S*-[3-(methoxycarbonyl)phenyl]-5’-thioadenosine (0.4 mmol) was stirred in dioxane (4 mL), forming an emulsion. LiAlH_4_ (2.4 mmol) was added, and the mixture was stirred at 50 °C for 3 hours.

The reaction mixture was neutralized by the addition of ethyl acetate (4 mL) and HCl (1 M, 2 mL), and then extracted into water (2 mL) three times. The grey solid was filtered out of the aqueous layer, washed with water, and dried *in vacuo* to afford the desired product as a clear, colorless oil in 44% yield. ^1^H (500 MHz; DMSO-d_6_): δ 8.36 (s, 1H), 8.14 (m, 1H), 7.25 (m, 5H), 7.12 (m, 1H), 5.89 (m, 1H), 5.62 (br s, 1H), 5.48 (br s, 1H), 5.31 (br s, 1H), 4.80 (m, 1H), 4.44 (s, 2H), 4.20 (m, 1H), 4.01 (m, 1H). ^13^C (125 MHz; DMSO-d_6_): δ 156.5 (C), 153.1(CH), 149.5 (C), 144.1 (C), 140.4 (CH), 135.9 (C), 129.2 (CH), 126.8 (CH), 126.5 (CH), 124.4 (CH), 119.6 (C), 87.9 (CH), 83.3 (CH), 73.1 (CH), 73.1 (CH), 62.9 (CH_2_), 35.7 (CH_2_); IR (neat, cm^-1^) 3350 (s), 2933 (w), 2833 (w), 2254 (w), 2128 (w), 1646 (s), 1599 (m), 1476 (m), 1422 (m), 1334 (m), 1236 (m), 1049 (s), 1023 (s), 996 (s), 825 (s), 764 (s), 572 (s); HRMS (ESI) calculated for [C_17_H_19_N_5_O_4_S + H^+^] 390.1231, found 390.1230.

*5’-S-[4-(hydroxymethyl)phenyl]-5’-thioadenosine (**6**).* Under an argon atmosphere, 4-mercaptobenzyl alcohol (0.314 mmol) was dissolved in DMF (5 mL). NaH (0.408 mmol) was added, and the resulting mixture was stirred for 30 minutes, becoming brown in color. 5’-*O*-Tosyladenosine (0.308 mmol) was added, and the mixture was stirred for a further 16 hours at 35 °C to give a light yellow-brown solution. The product was concentrated *in vacuo*, dissolved in 1:1 water:acetonitrile (10 mL) and HCl (1 M, 0.5 mL), concentrated *in vacuo* to 1 mL, and then recrystallized at 0 °C over 16 hours. The obtained crystals were filtered with cold water, dried, and triturated with ethanol to afford the desired product as a clear, colorless oil in 18% yield. ^1^H (500 MHz; MeOD): δ 8.08 (s, 1H), 8.08 (s, 1H), 7.26 (m, 2H), 7.13 (m, 2H), 5.86 (m, 1H), 4.44 (s, 2H), 4.24 (m, 1H), 4.11 (m, 1H), 3.22 (m, 1H), 3.17(m, 1H); ^13^C (125 MHz; MeOD): δ 157.3 (C), 153.9 (CH), 150.7 (C), 141.4 (C), 141.2 (CH), 136.0 (C), 130.7 (CH), 128.7 (CH), 120.6 (C), 90.1 (CH), 85.1 (CH), 74.8 (CH), 74.1 (CH), 64.7 (CH_2_), 37.3 (CH_2_); IR (neat, cm^-1^) 3316 (m), 3177 (m), 2899 (m), 1645 (s), 1599 (s), 1576 (s), 1478 (m), 1418 (m), 1334 (m), 1295 (m), 1212 (m), 1125 (s), 1088 (s), 1035 (s), 796 (s), 642 (s); HRMS (ESI) calculated for [C_17_H_19_N_5_O_4_S + H^+^] 390.1231, found 390.1231.

### Synthesis of PTA derivatives 9–12

5*’*-*S*-(3-acetaminophenyl)-5*’*-thioadenosine *(**9**).* 3-Acetamidothiophenol (0.788 mmol) was dissolved in dry DMF (10 mL) and placed under an argon atmosphere. Potassium *tert*-butoxide (1M in THF; 0.866 mmol) was added, and the mixture was stirred at room temperature for 30 minutes, after which 5*’*-chloro-5*’*-deoxyadenosine (0.525 mmol) was added. Stirring was continued at 40 °C for 16 hours, after which the mixture was neutralized with 10% aqueous HCl, and concentrated *in vacuo*. The crude product was recrystallized from water (2 mL) by heating to 60 °C and then cooling at 4 °C for 16 hours. Obtained crystals were rinsed with cold water followed by dichloromethane, to afford the desired product as white crystalline solid in 88% yield. ^1^H (500 MHz; DMSO-d_6_): δ 9.95 (s, 1H), 8.34 (s, 1H), 8.15 (s, 1H), 7.62 (m, 1H), 7.37 (m, 1H), 7.27 (br s, 2H), 7.21 (m, 1H), 7.01 (m, 1H), 5.87 (d, *J* = 6.0 Hz, 1H), 5.58 (br s, 1H), 5.44 (br s, 1H), 4.81 (m, 1H), 4.19 (m, 1H), 4.00 (m, 1H), 3.38 (dd, 1H), 3.26 (dd, *J* = 13.8, 7.0 Hz, 1H); ^13^C (125 MHz; DMSO-d_6_): δ 168.5 (C), 156.0 (C), 152.6 (CH), 149.4 (C), 139.9 (CH), 139.8 (C), 136.1 (C), 129.3 (CH), 122.5 (CH), 119.1 (C), 118.2 (CH), 116.5 (CH), 87.4 (CH), 82.7 (CH), 72.6 (CH), 72.5 (CH), 35.1 (CH_2_), 24.0 (CH_3_); IR (neat, cm^-1^) 3267 (m), 3127 (m), 2886 (w), 1652 (s), 1605 (s), 1587 (s), 1544 (s), 1477 (s), 1420 (s), 1295 (s), 1206 (m), 1090 (s), 1044 (s), 776 (m), 684 (s), 638 (s), 537 (s); HRMS (ESI) calculated for [C_18_H_20_N_6_O_4_S + H^+^] 417.1340, found 417.1340; M.P. 83–86 °C.

5*’*-*S*-(4-acetaminophenyl)-5*’*-thioadenosine *(**10**).* 4-Acetamidothiophenol (1.05 mmol) was dissolved in dry DMF (10 mL) and placed under an argon atmosphere. Potassium *tert*-butoxide (1M in THF; 1.16 mmol) was added, and the mixture was stirred at room temperature for 30 minutes, after which 5*’*-chloro-5*’*-deoxyadenosine (0.525 mmol) was added. Stirring was continued at 40 °C for 16 hours, after which the mixture was neutralized with 10% aqueous HCl, and concentrated *in vacuo*. The crude product was recrystallized from 7 mL of 5:2 water:methanol by heating to 70 °C and then cooling at 4 °C for 16 hours. Obtained crystals were rinsed with cold water followed by dichloromethane, to afford the desired product as white crystalline solid in 57% yield. ^1^H (500 MHz; DMSO-d_6_; TFA salt): δ 9.96 (s, 1H), 8.78 (s, 1H), 8.47 (s, 1H), 7.53 (m, 2H), 7.31 (m, 2H), 7.28 (br s, 2H), 5.88 (d, *J* = 6.0 Hz, 1H), 5.52 (br d, 1H), 5.36 (br d, 1H), 4.79 (m, 1H), 4.16 (m, 1H), 3.69 (m, 1H), 3.34 (dd, *J* = 13.9, 6.2 Hz, 1H), 3.21 (dd, *J* = 13.8, 7.1 Hz, 1H), 2.02 (s, 3H); ^13^C (125 MHz; DMSO-d_6_, TFA salt): δ 168.2 (C), 156.0 (C), 152.6 (CH), 149.4 (C), 139.9 (CH), 137.9 (C), 130.1 (CH), 128.7 (C), 119.6 (CH), 119.2 (C), 87.3 (CH), 82.8 (CH), 72.6 (CH), 72.5 (CH), 36.4 (CH2), 23.9 (CH_3_); IR (neat, cm^-1^) 3266 (m), 3111 (m), 2879 (w), 1646 (s), 1593 (s), 1533 (s), 1495 (s), 1396 (s), 1295 (s), 1088 (s), 1042 (s), 815 (s), 716 (s), 648 (s), 503 (s); HRMS (ESI) calculated for [C_18_H_20_N_6_O_4_S + Na^+^] 439.1159, found 439.1157; M.P. 94–96 °C.

*5’-S-[3-[(aminocarbonyl)amino]phenyl]-5’-thioadenosine (**11**).* m-APTA (0.400 mmol) was ground into methanol (8 mL) and glacial acetic acid (0.5 mL), while stirring at 0 °C. KOCN (1.60 mmol) was dissolved in water (1 mL) and transferred to the stirring solution. The reaction mixture was warmed to 20 °C with stirring over 30 minutes, then heated to 50 °C with continued stirring for 4 hours. The mixture was then neutralized by addition of NaOH (10 wt%, 4.5 mL) and concentrated *in vacuo* to afford a cloudy aqueous mixture. This was crystallized at 4 °C to afford the desired product as a white solid in 40% yield. ^1^H (500 MHz; DMSO-d_6_): δ 9.09 (s, 1H), 8.34 (s, 1H), 8.15 (s, 1H), 7.47 (m, 1H), 7.26 (br s, 2H), 7.22 (m, 1H), 7.12 (m, 1H), 6.86 (m, 1H), 6.06 (br s, 2H), 5.88 (d, *J* = 6.0 Hz, 1H), 4.78 (m, 1H), 4.18 (m, 1H), 4.01 (m, 1H), 3.37 (dd, 1H), 3.23 (dd, *J* = 13.8, 7.1 Hz, 1H); ^13^C (125 MHz; DMSO-d_6_): δ 156.1 (C), 156.0 (C), 153.1 (C), 152.6 (CH), 141.4 (C), 139.9 (CH), 135.8 (C), 129.1 (CH), 120.4 (CH), 119.1 (C), 117.0 (CH),115.3 (CH), 87.4 (CH), 82.8 (CH), 72.6 (CH), 72.6 (CH), 35.2 (CH_2_); IR (neat, cm^-1^) 3304 (m), 3177 (m), 2893 (w), 1652 (s), 1576 (s), 1544 (s), 1476 (s), 1333 (s), 1300 (s), 1245 (s), 1207 (s), 1090 (s), 1045 (s), 686 (s), 646 (s); HRMS (ESI) calculated for [C_17_H_19_N_7_O_4_S + Na^+^] 440.1111, found 440.1111; M.P. 156–157 °C.

*5’-S-[4-[(aminocarbonyl)amino]phenyl]-5’-thioadenosine (**12**).* p-APTA (0.400 mmol) was ground into methanol (8 mL) and glacial acetic acid (0.5 mL), while stirring at 0 °C. KOCN (1.60 mmol) was dissolved in water (1 mL) and transferred to the stirring mixture. The reaction mixture was warmed to 20 °C with stirring over 30 minutes, then heated to 50 °C with continued stirring for 4 hours. The mixture was then neutralized by addition of NaOH (10 wt%, 4.7 mL) and concentrated *in vacuo* to afford a cloudy aqueous mixture. This was crystallized at 4 °C to afford the desired product as a white solid in 75% yield. ^1^H (500 MHz; DMSO-d_6_; TFA salt): δ 8.64 (s, 1H), 8.62 (s, 1H), 8.43 (s, 1H), 7.36 (m, 2H), 7.26 (m, 2H), 5.92 (d, *J* = 6.1 Hz, 1H), 4.70 (m, 1H), 4.14 (m, 1H), 3.98 (m, 1H), 3.28 (dd, *J* = 13.9, 6.3 Hz, 1H), 3.15 (dd, *J* = 13.9, 6.9 Hz, 1H); ^13^C (125 MHz; DMSO-d_6_, TFA salt): δ 155.9 (C), 155.8 (C), 151.6 (CH), 148.7 (C), 147.0 (C), 141.9 (C), 139.6 (CH), 131.1 (CH), 125.8 (C), 118.4 (CH), 87.5 (CH), 83.4 (CH), 73.1 (CH), 72.4 (CH), 37.0 (CH_2_); IR (neat, cm^-1^) 3311 (m), 3113 (m), 2893 (w), 1674 (s), 1576 (s), 1534 (s), 1496 (s), 1402 (s), 1342 (s),1297 (s), 1211 (s), 1130 (s), 1087 (s), 1045 (s), 817 (s), 719 (s), 639 (s); HRMS (ESI) calculated for [C_17_H_19_N_7_O_4_S + Na^+^] 440.1111, found 440.1110; M.P. 141–144 °C.

## Spectral Data for Synthetic Materials

**Fig. S7.**
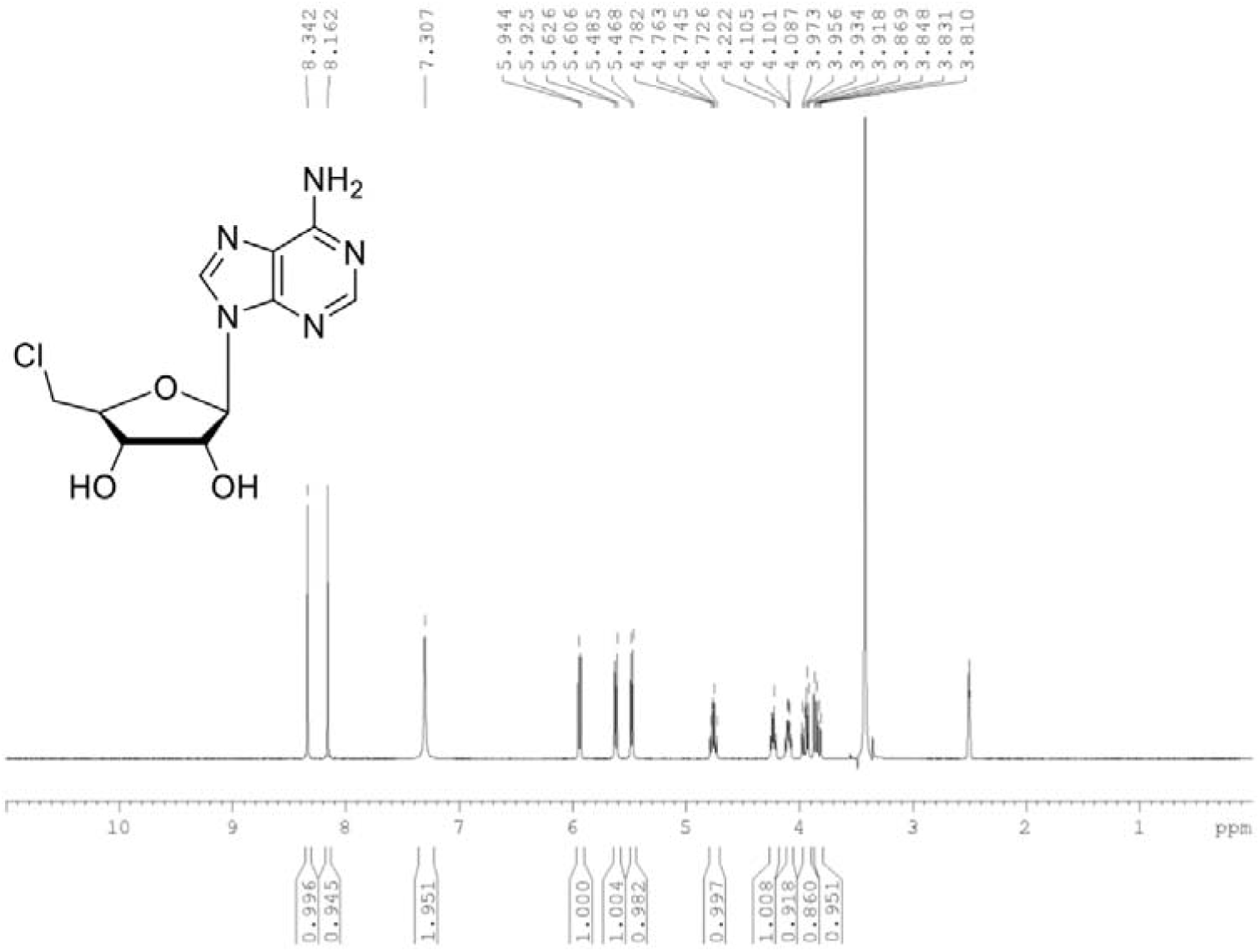
^1^H NMR spectrum for 5*’*-chloro-5*’*-deoxyadenosine in DMSO-d_6_ at 300 MHz

**Fig. S8.**
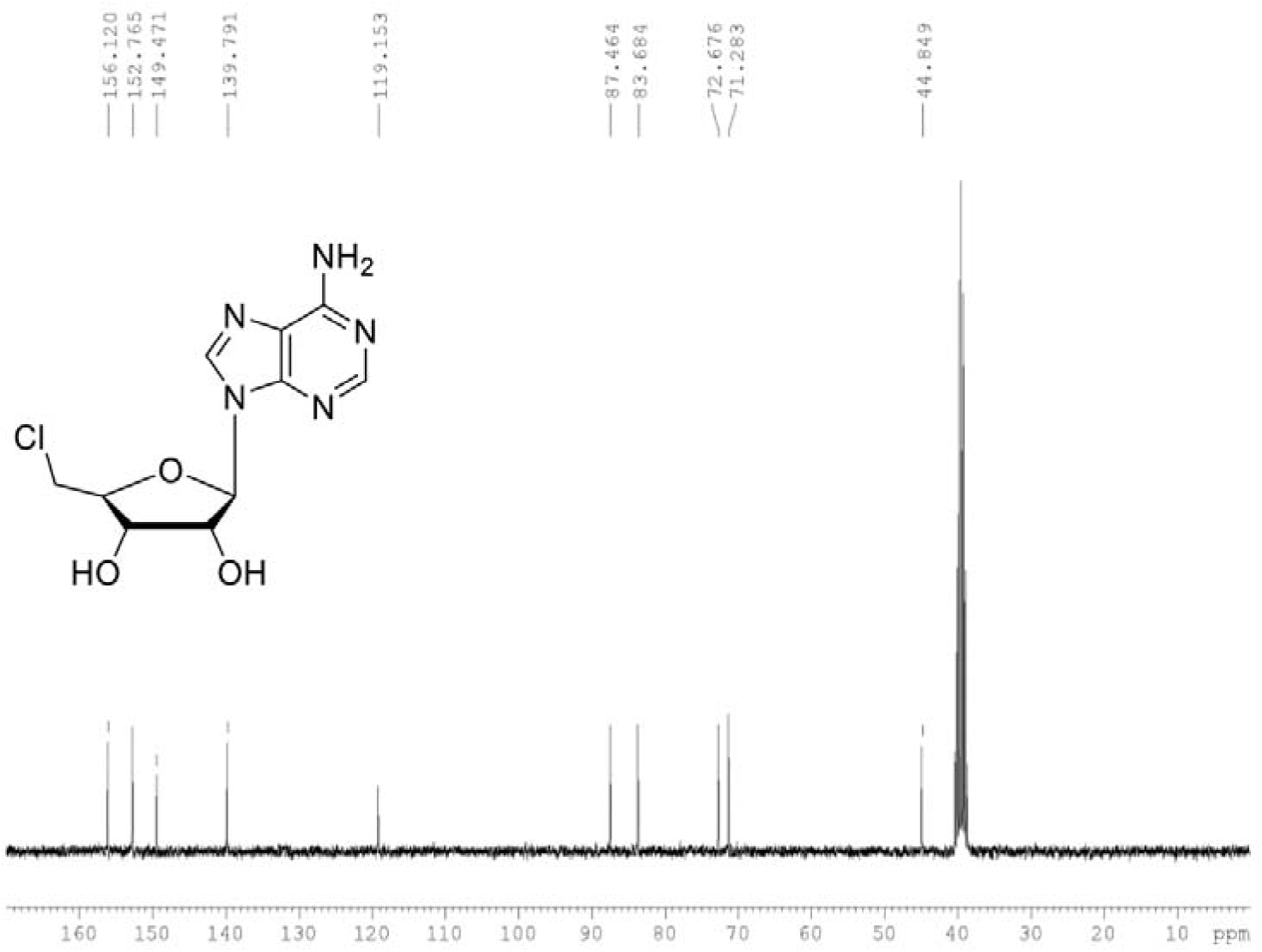
^13^C NMR spectrum for 5*’*-chloro-5*’*-deoxyadenosine in DMSO-d_6_ at 75 MHz

**Fig. S9.**
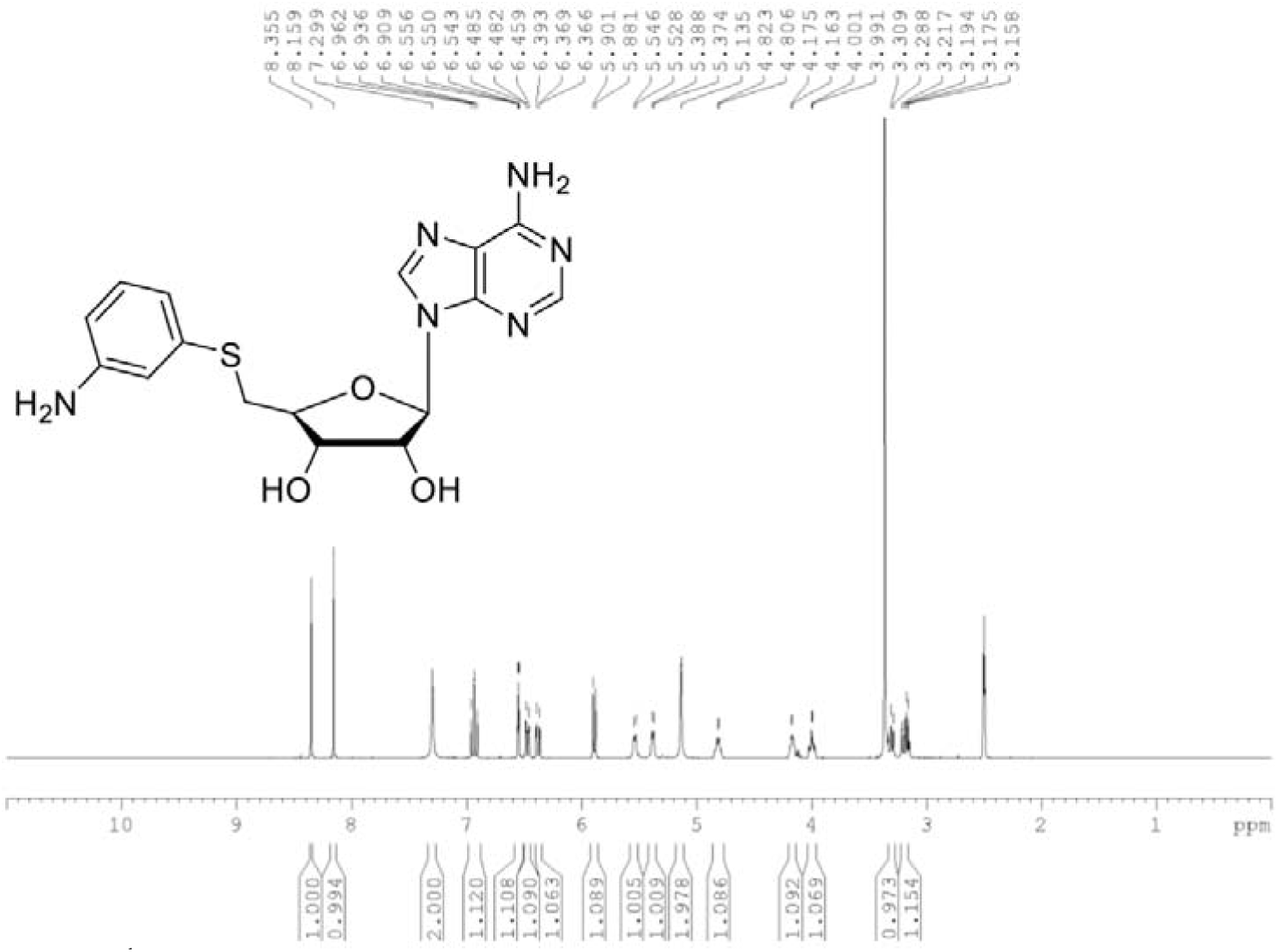
^1^H NMR spectrum for m-APTA in DMSO-d_6_ at 300 MHz

**Fig. S10.**
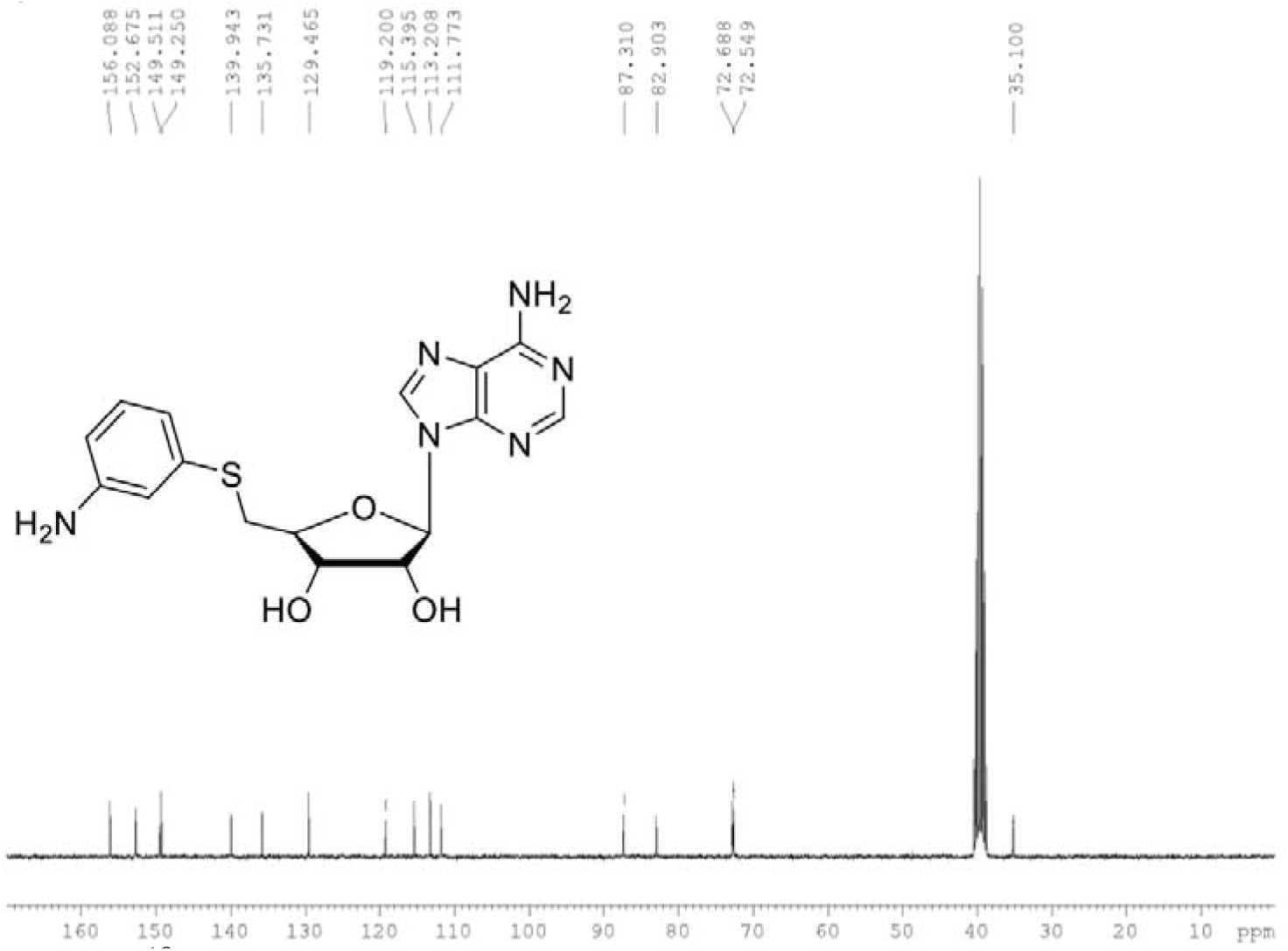
^13^C NMR spectrum for m-APTA in DMSO-d_6_ at 75 MHz

**Fig. S11.**
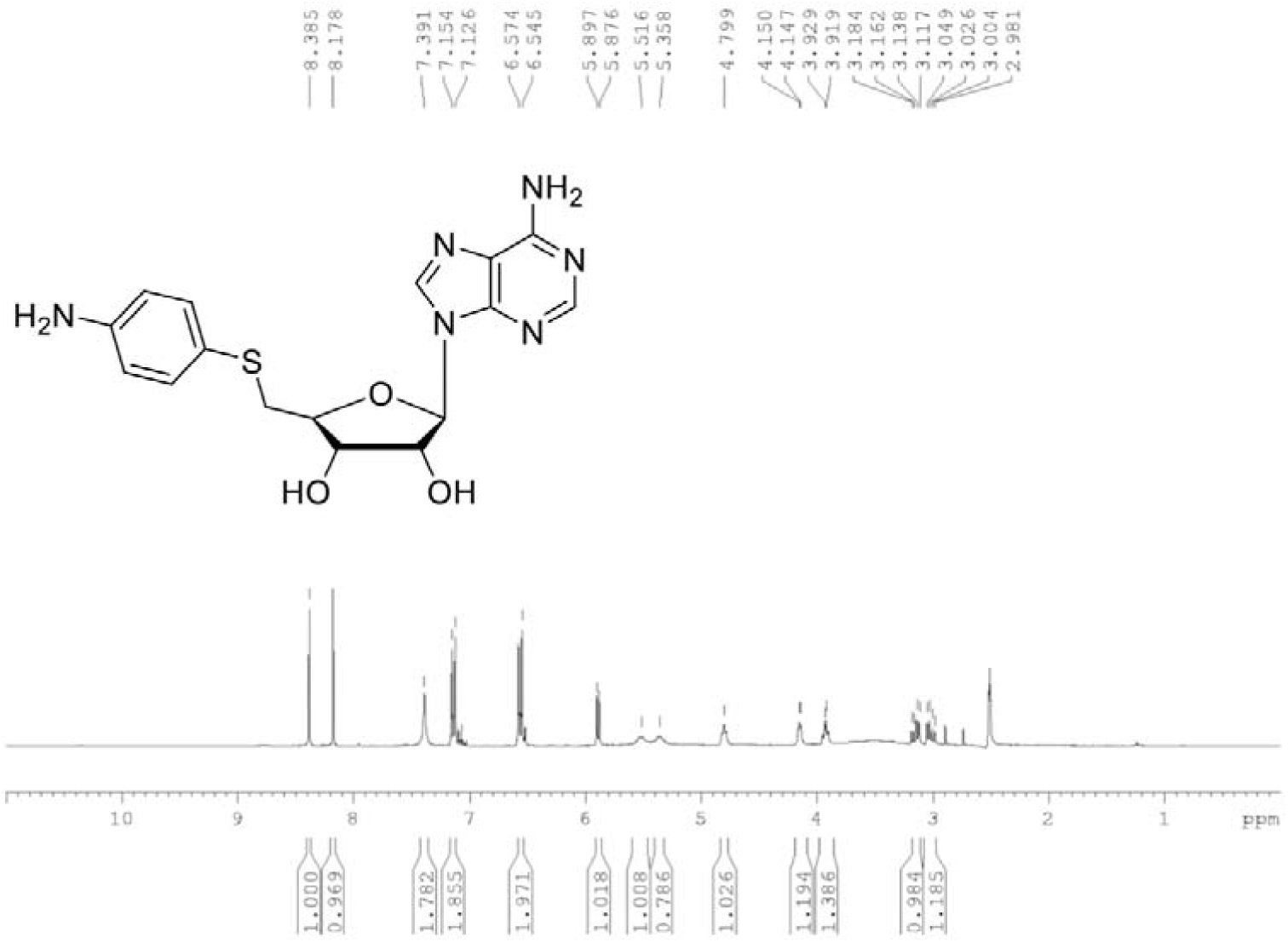
^1^H NMR spectrum for p-APTA in DMSO-d_6_ at 300 MHz

**Fig. S12.**
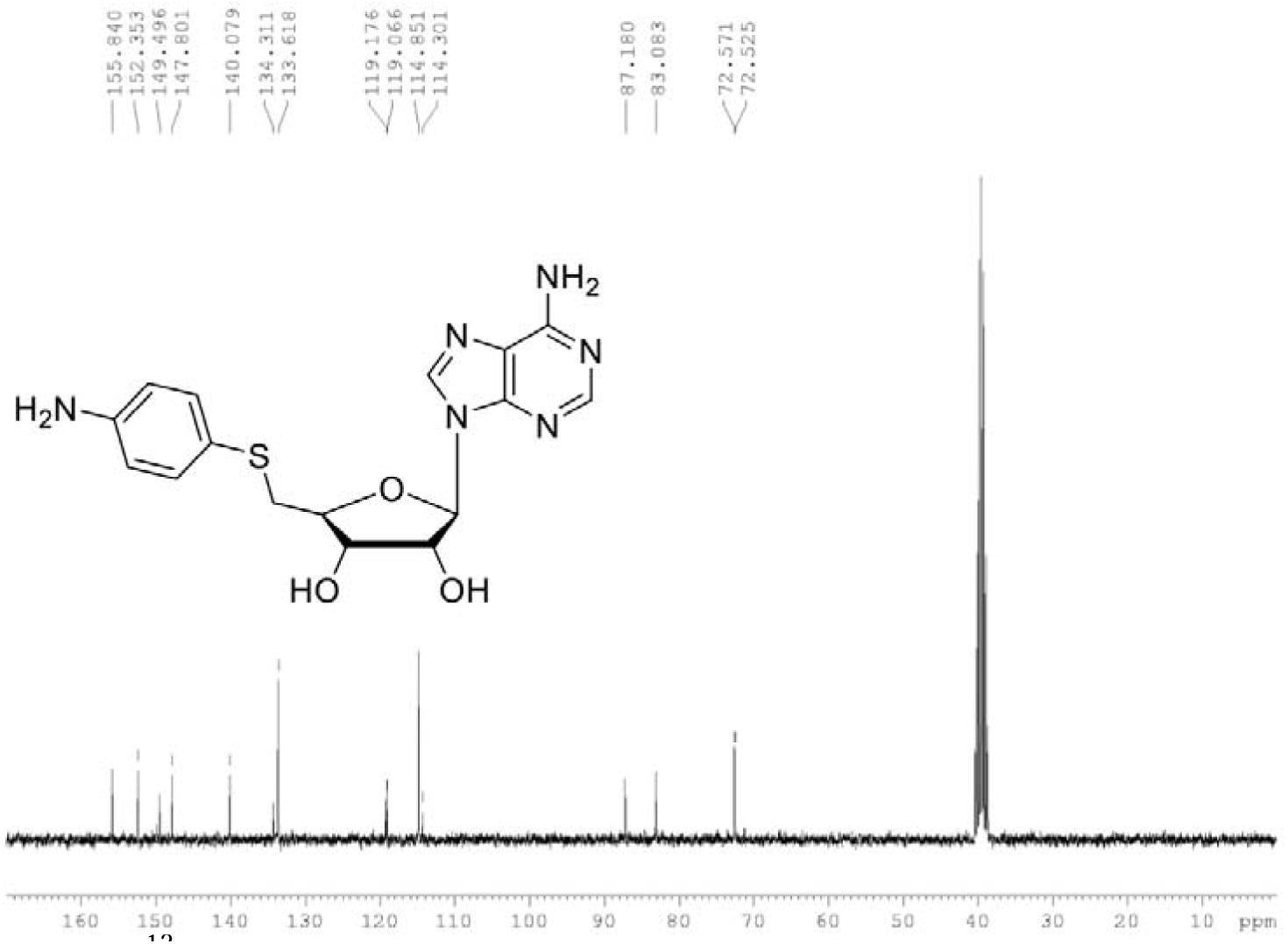
^13^C NMR spectrum for p-APTA in DMSO-d_6_ at 75 MHz

**Fig. S13.**
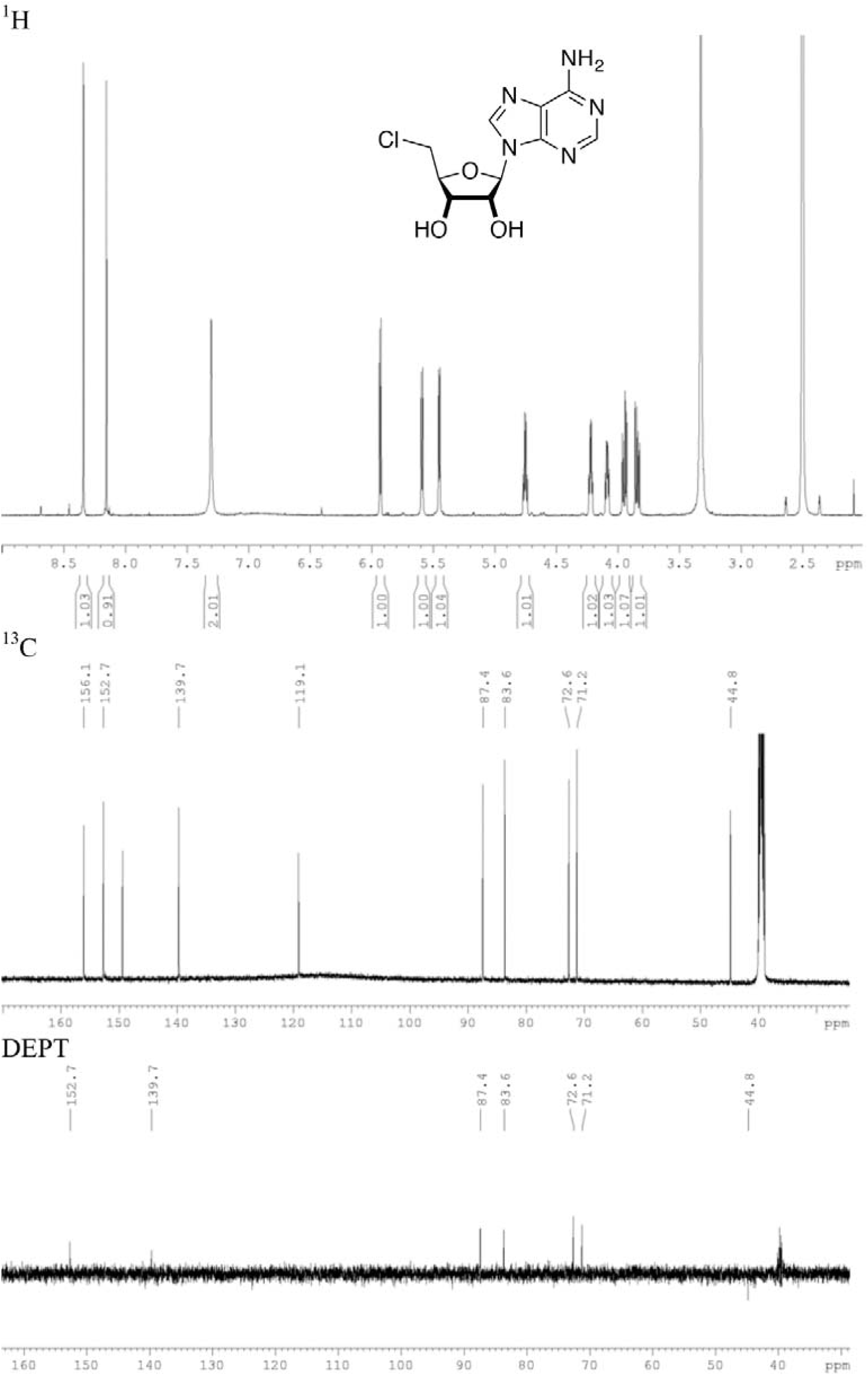
^1^H, ^13^C and DEPT-135 NMR spectra for 5*’*-chloro-5*’*-deoxyadenosine in DMSO-d_6_ at 500 and 125 MHz

**Fig. S14.**
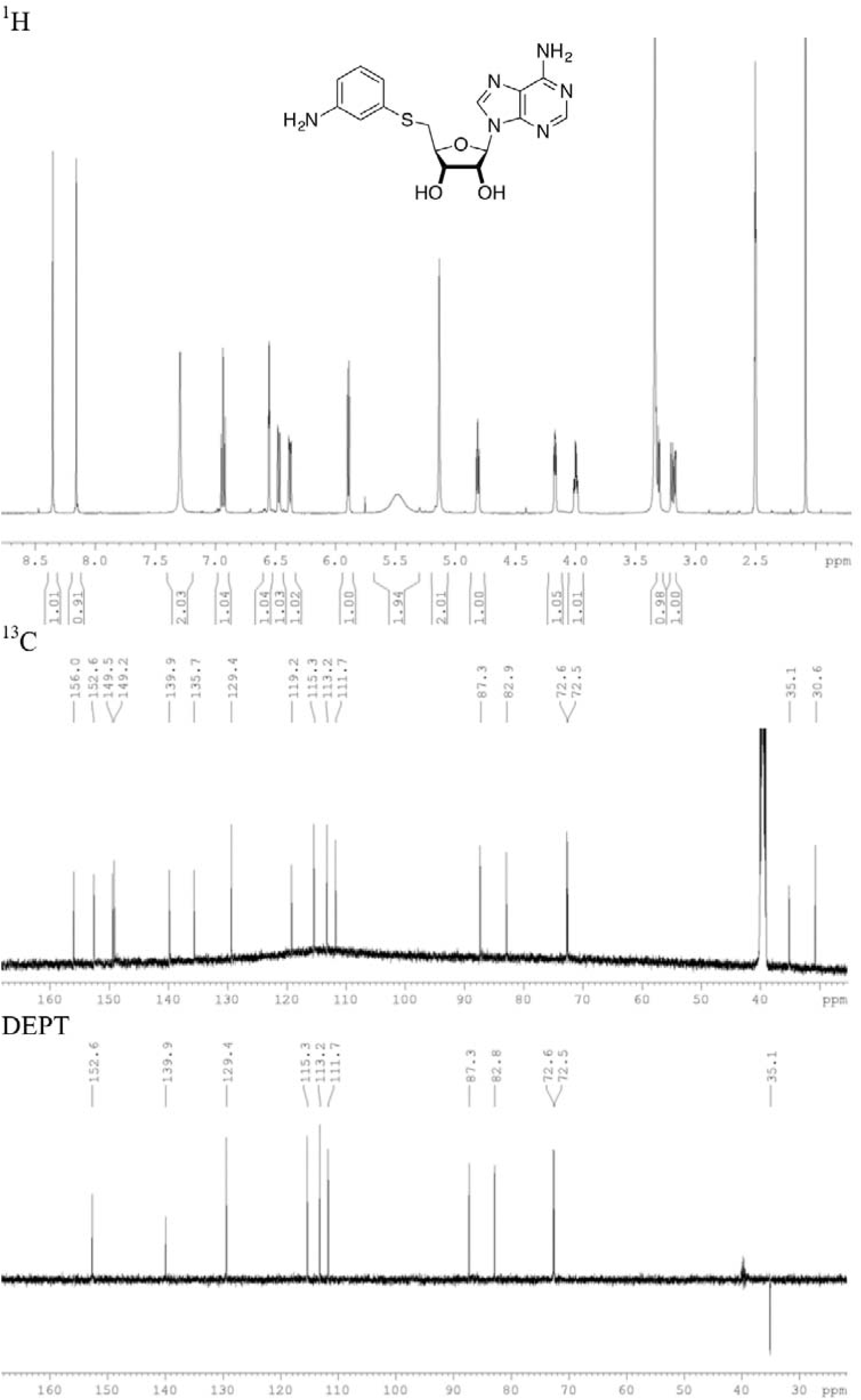
^1^H, ^13^C and DEPT-135 NMR spectra for m-APTA in DMSO-d_6_ at 500 and 125 MHz

**Fig. S15.**
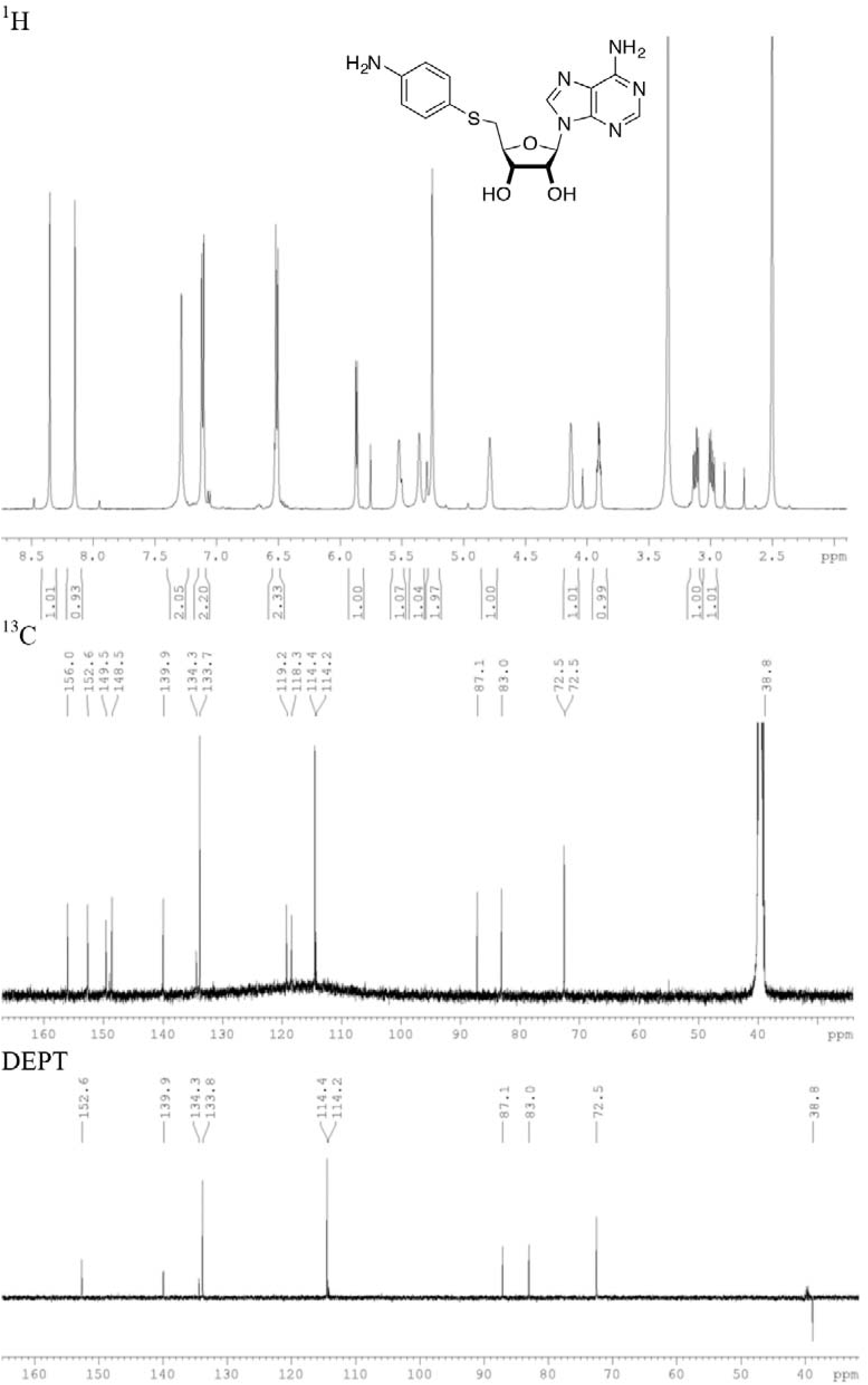
^1^H, ^13^C and DEPT-135 NMR spectra for p-APTA in DMSO-d_6_ at 500 and 125 MHz

**Fig. S16.**
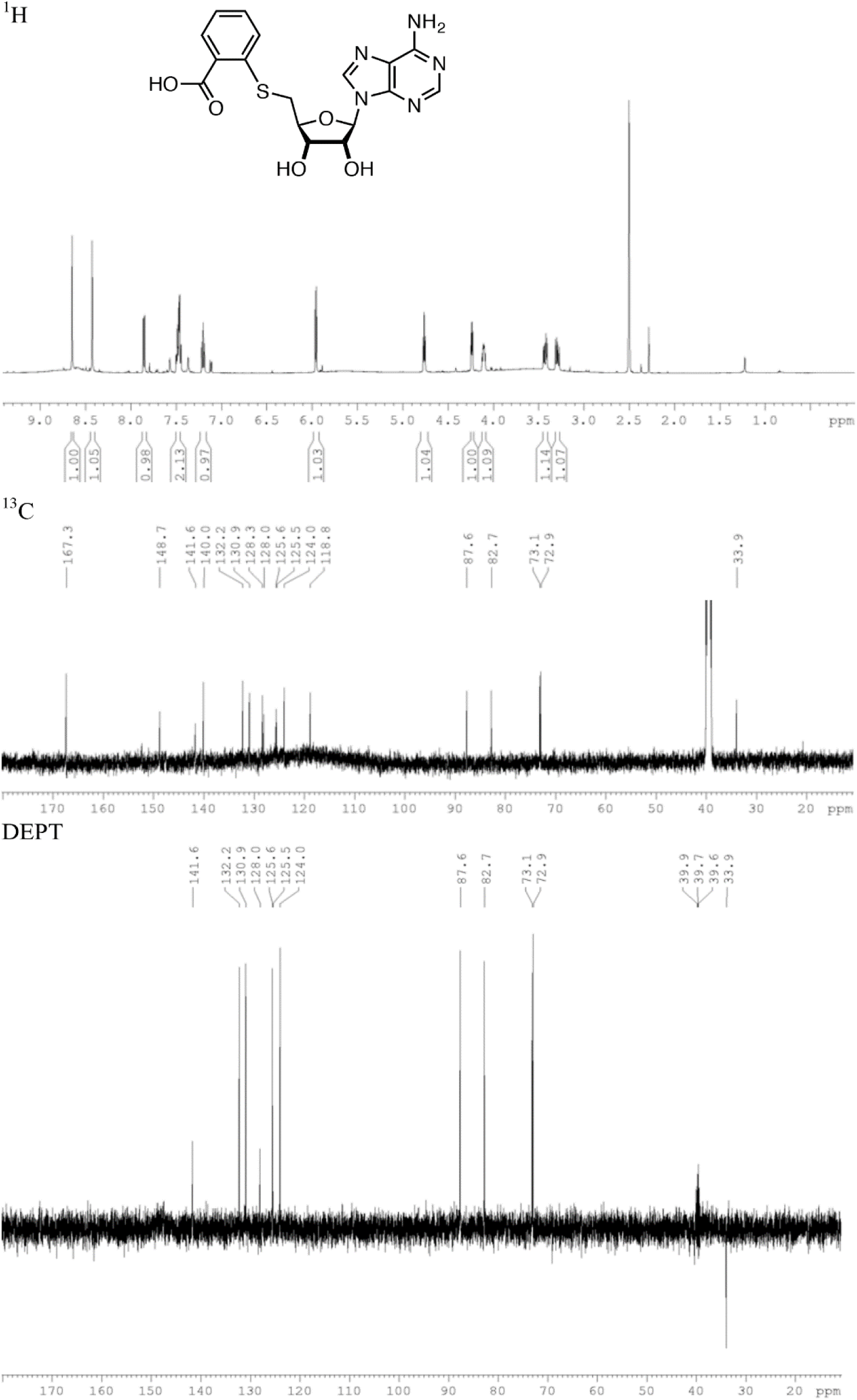
^1^H, ^13^C and DEPT-135 NMR spectra for derivative **1** in DMSO-d_6_ at 500 and 125 MHz

**Fig. S17.**
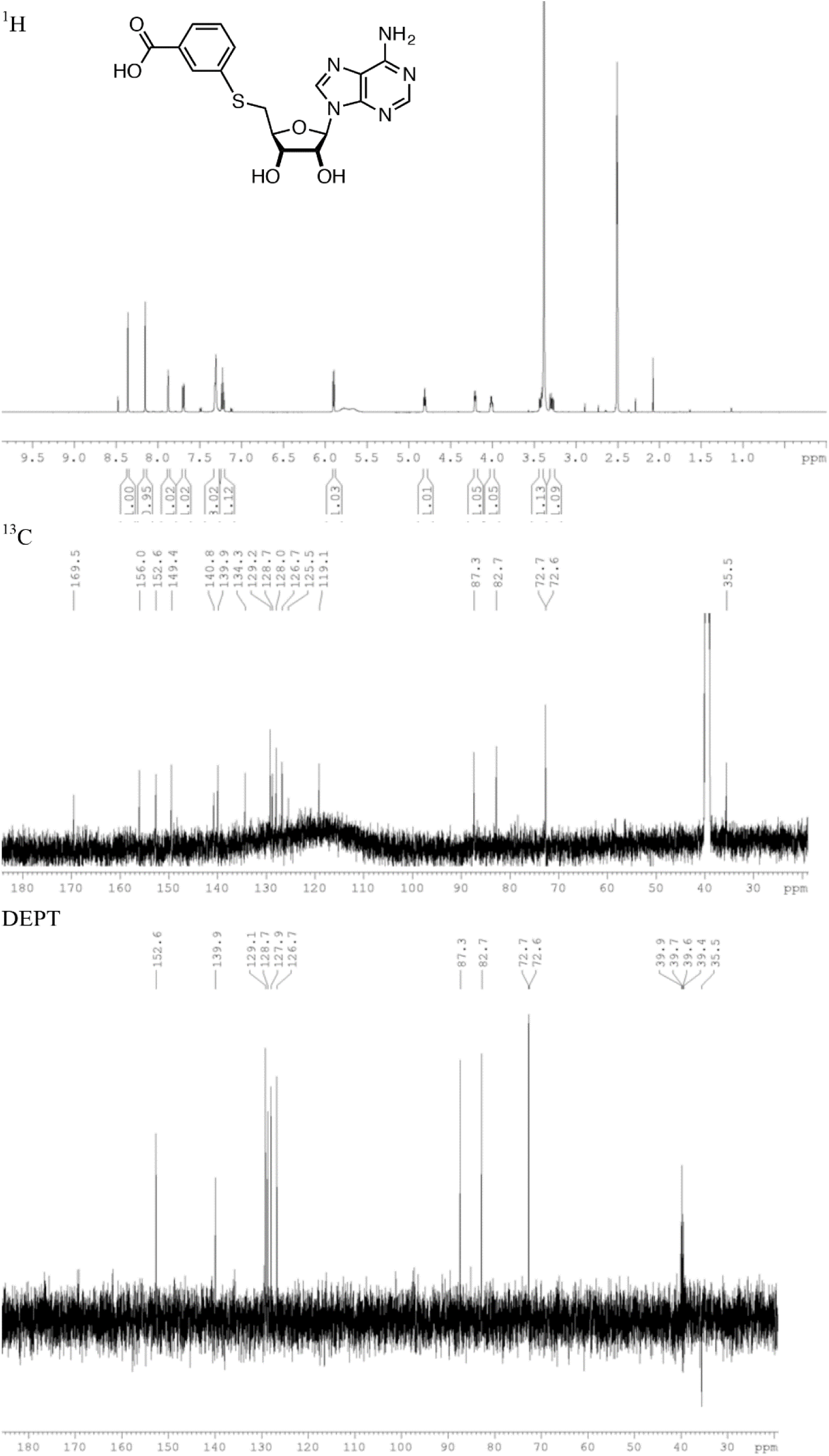
^1^H, ^13^C and DEPT-135 NMR spectra for derivative **2** in DMSO-d_6_ at 500 and 125 MHz

**Fig. S18.**
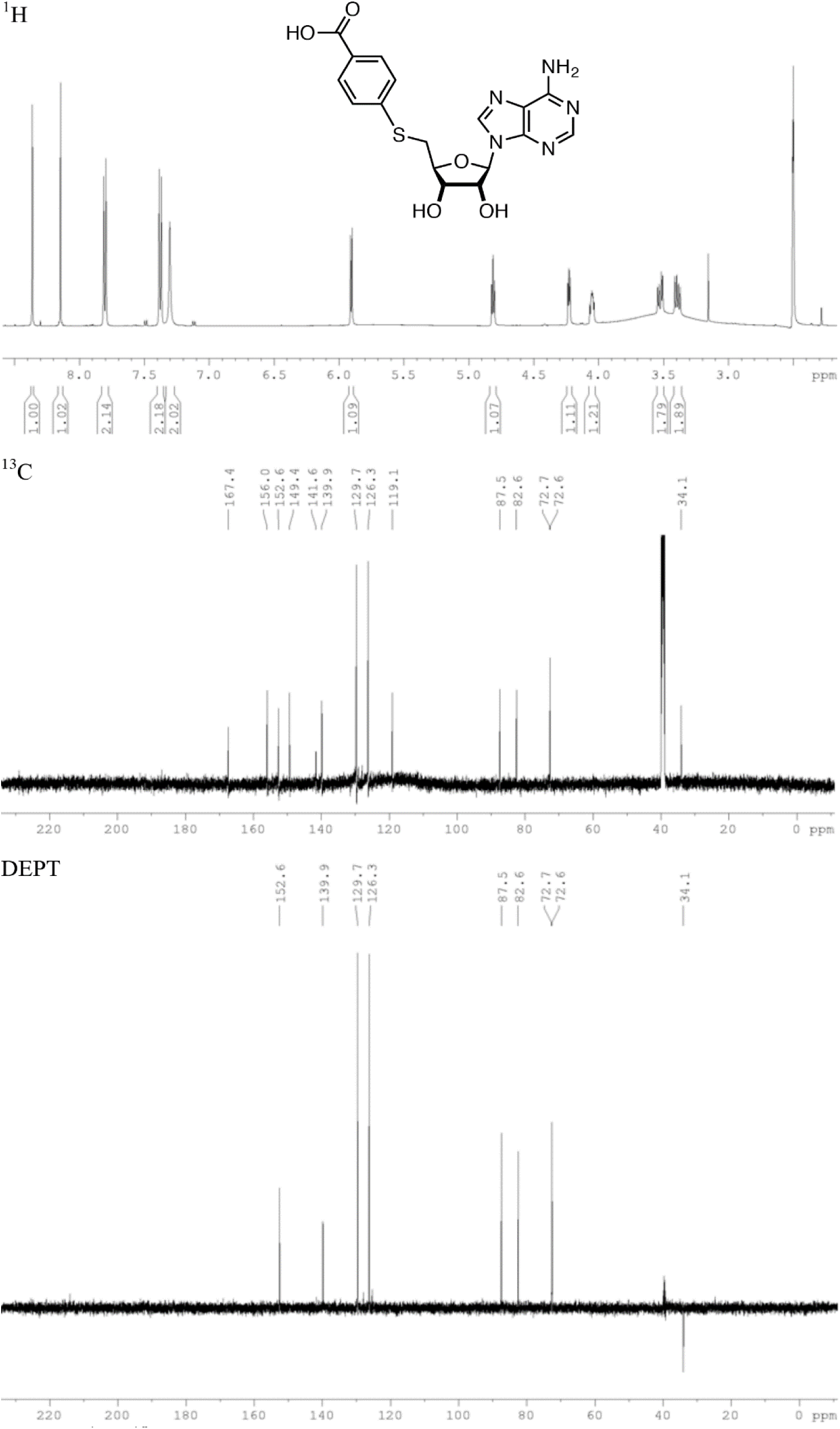
^1^H, ^13^C and DEPT-135 NMR spectra for derivative **3** in DMSO-d_6_ at 500 and 125 MHz

**Fig. S19.**
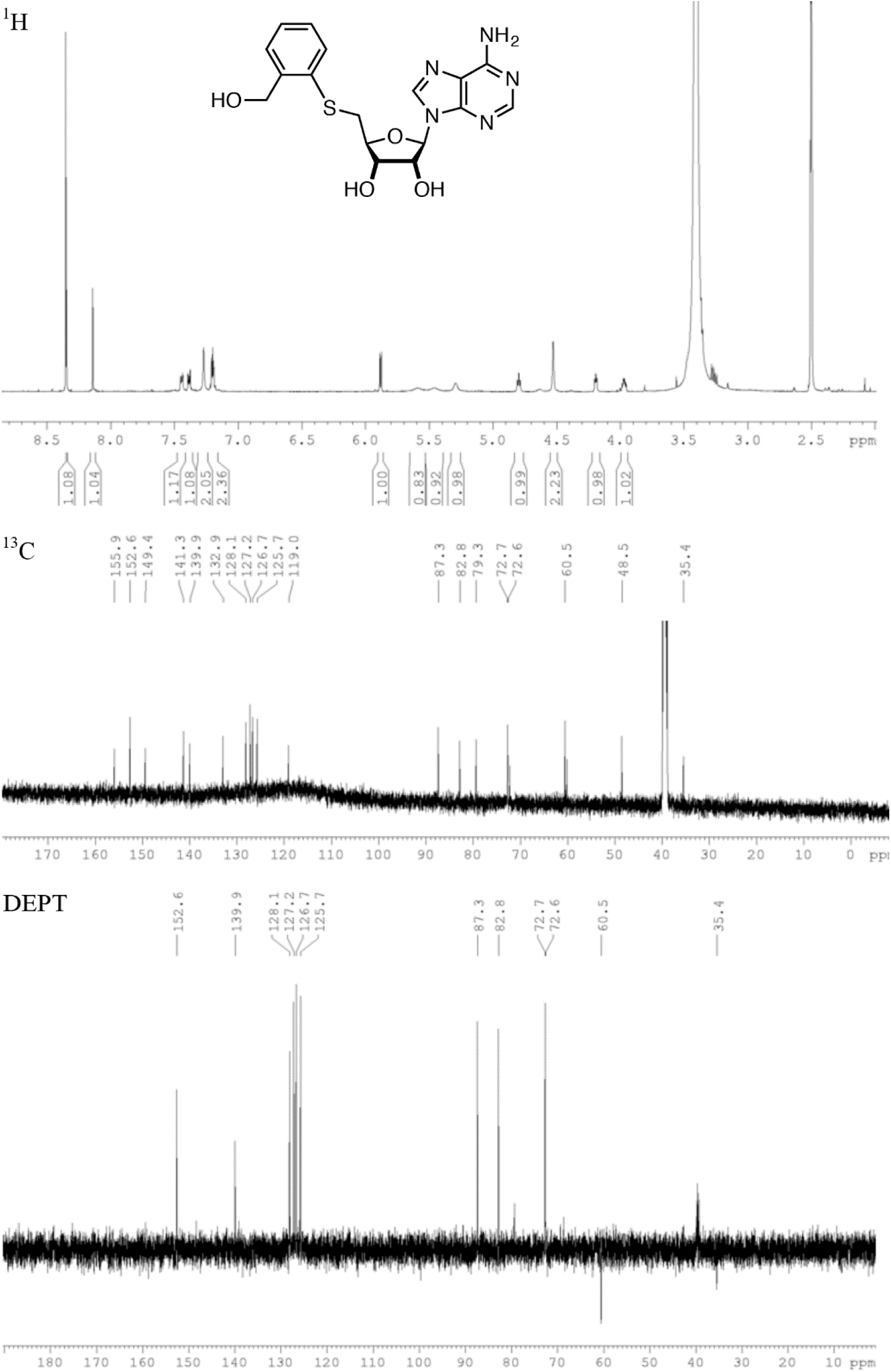
^1^H, ^13^C and DEPT-135 NMR spectra for derivative **4** in DMSO-d_6_ at 500 and 125 MHz

**Fig. S20.**
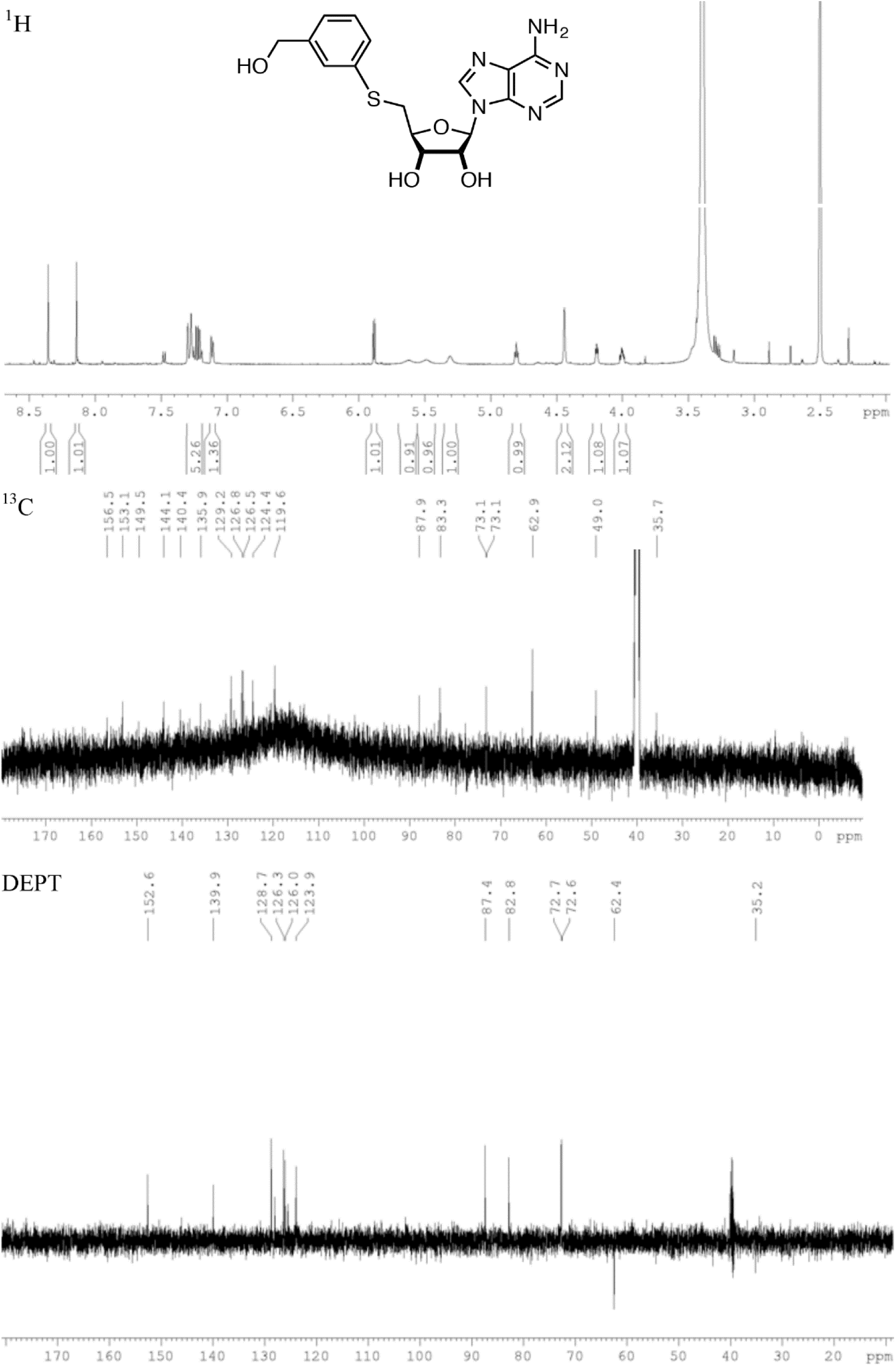
^1^H, ^13^C and DEPT-135 NMR spectra for derivative **5** in DMSO-d_6_ at 500 and 125 MHz

**Fig. S21.**
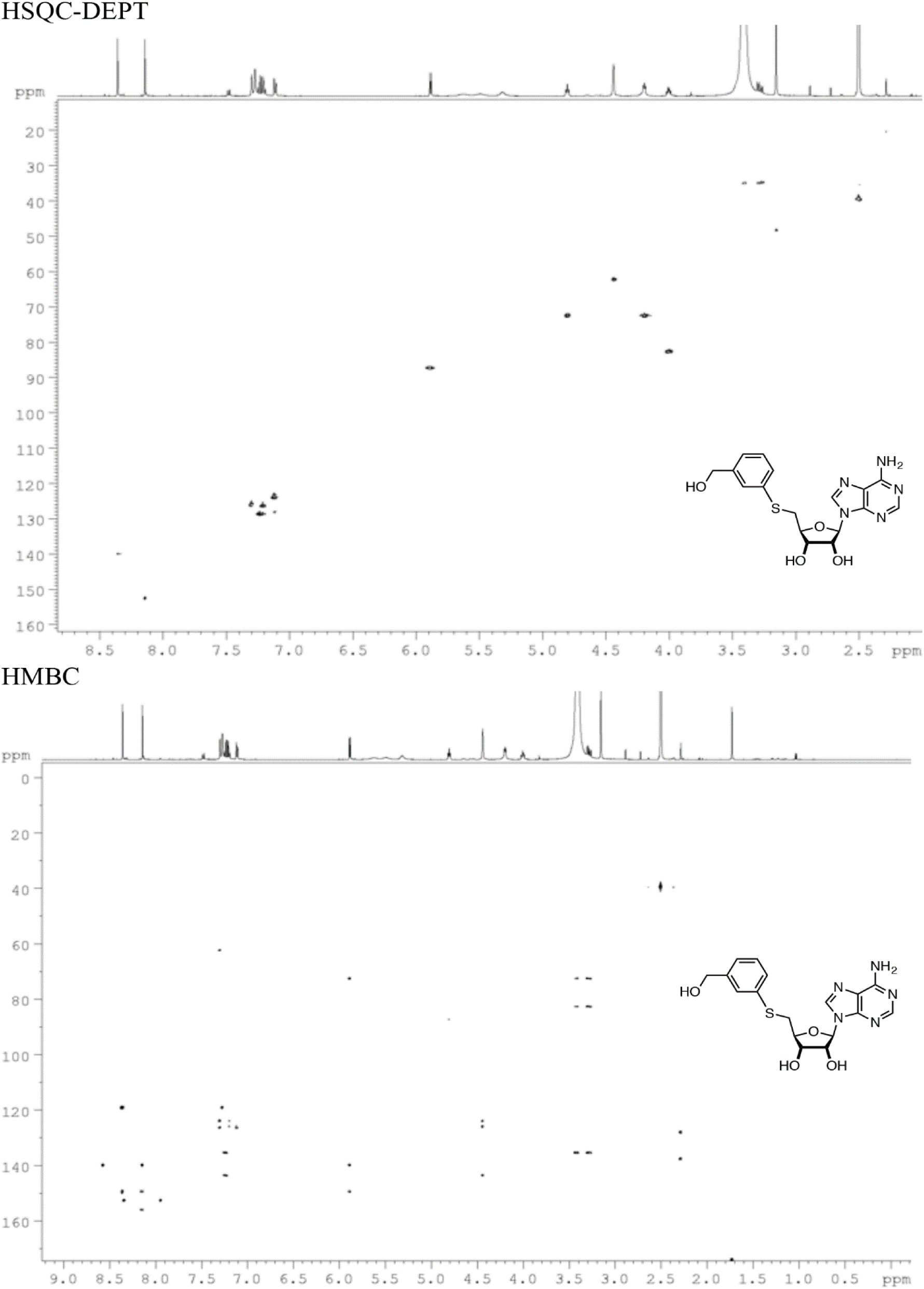
HSQC and HMBC spectra for derivative **5** in DMSO-d_6_

**Fig. S22.**
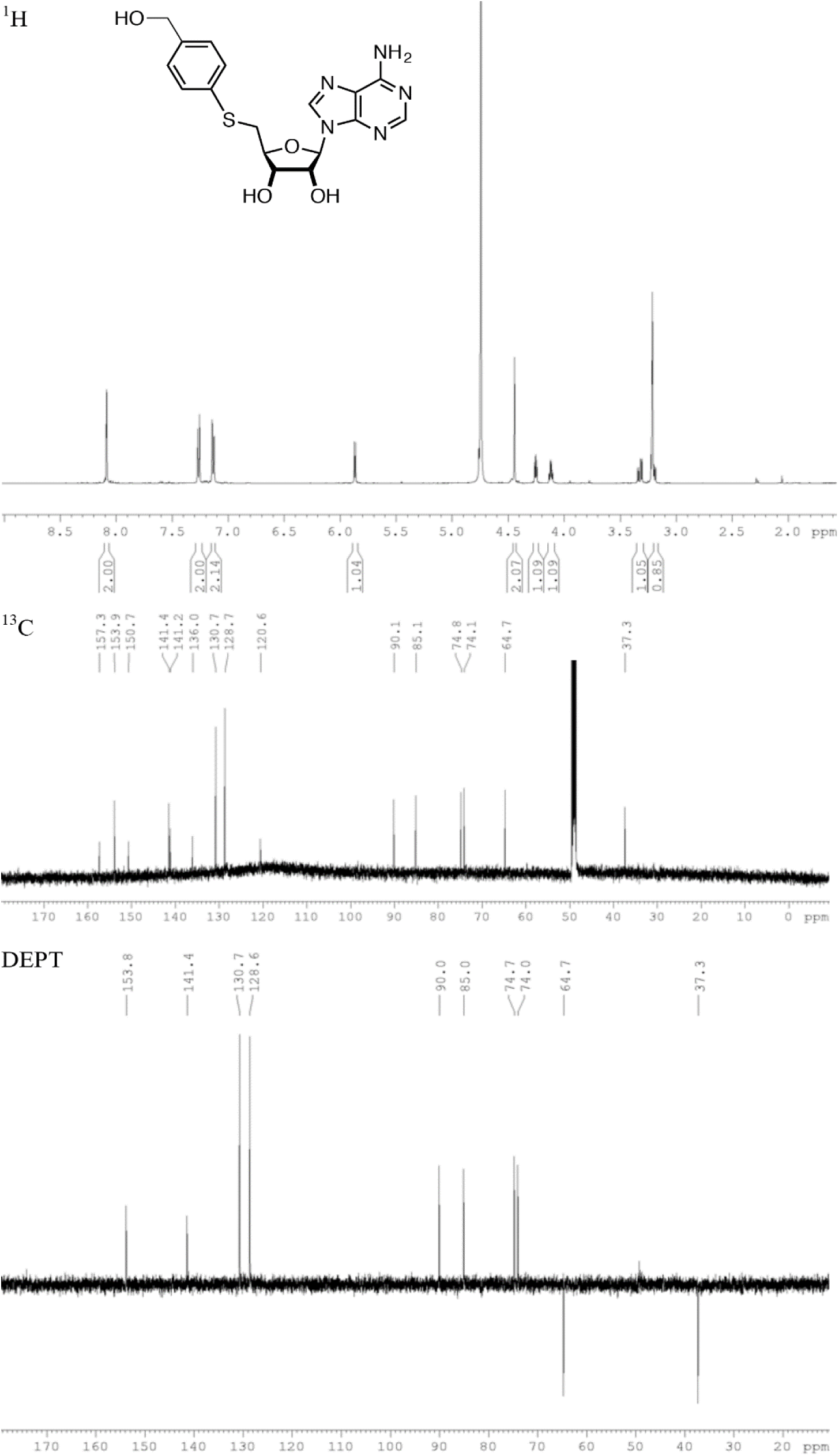
^1^H, ^13^C and DEPT-135 NMR spectra for derivative **6** in MeOD at 500 and 125 MHz

**Fig. S23.**
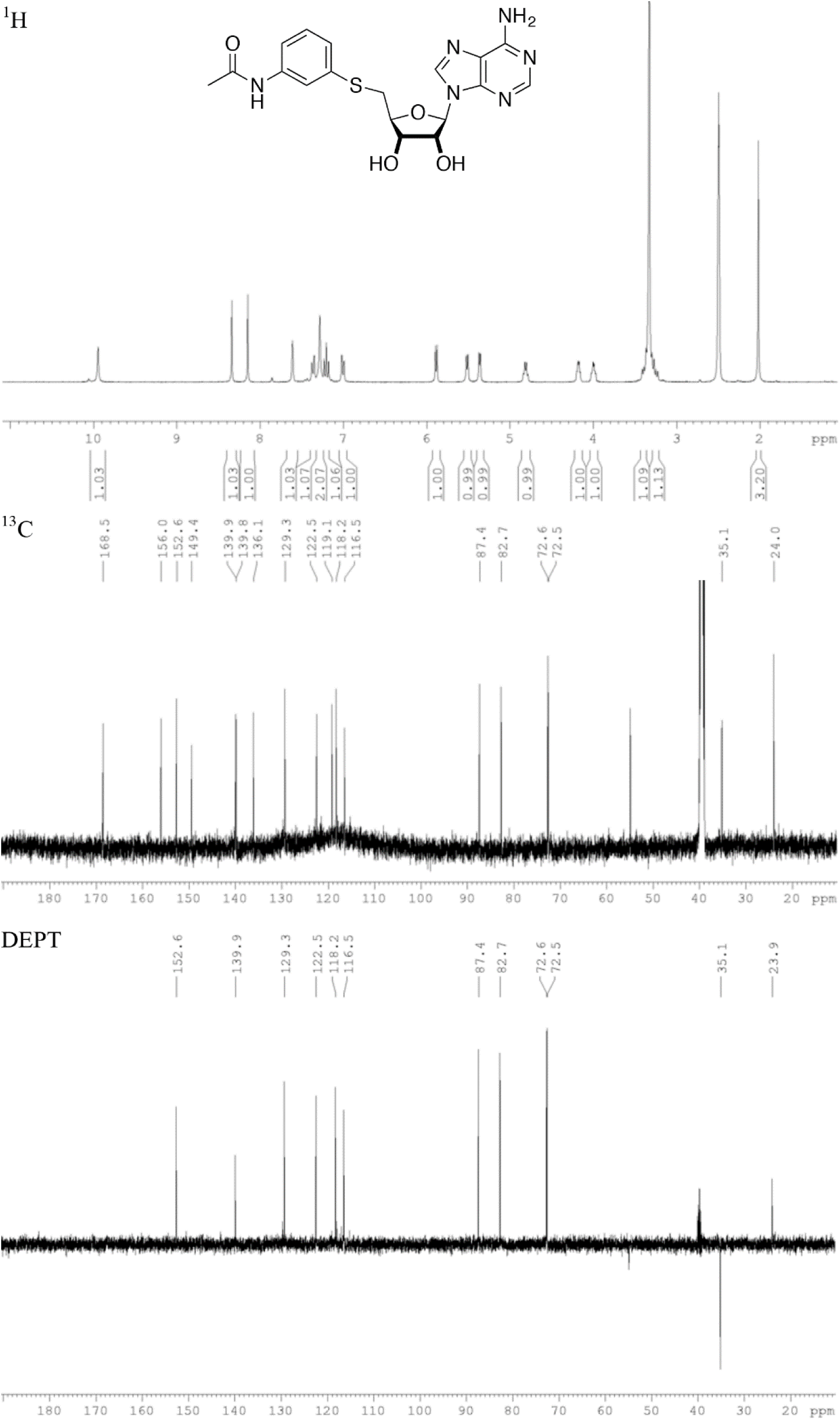
^1^H, ^13^C and DEPT-135 NMR spectra for derivative **9** in DMSO-d_6_ at 500 and 125 MHz

**Fig. S24.**
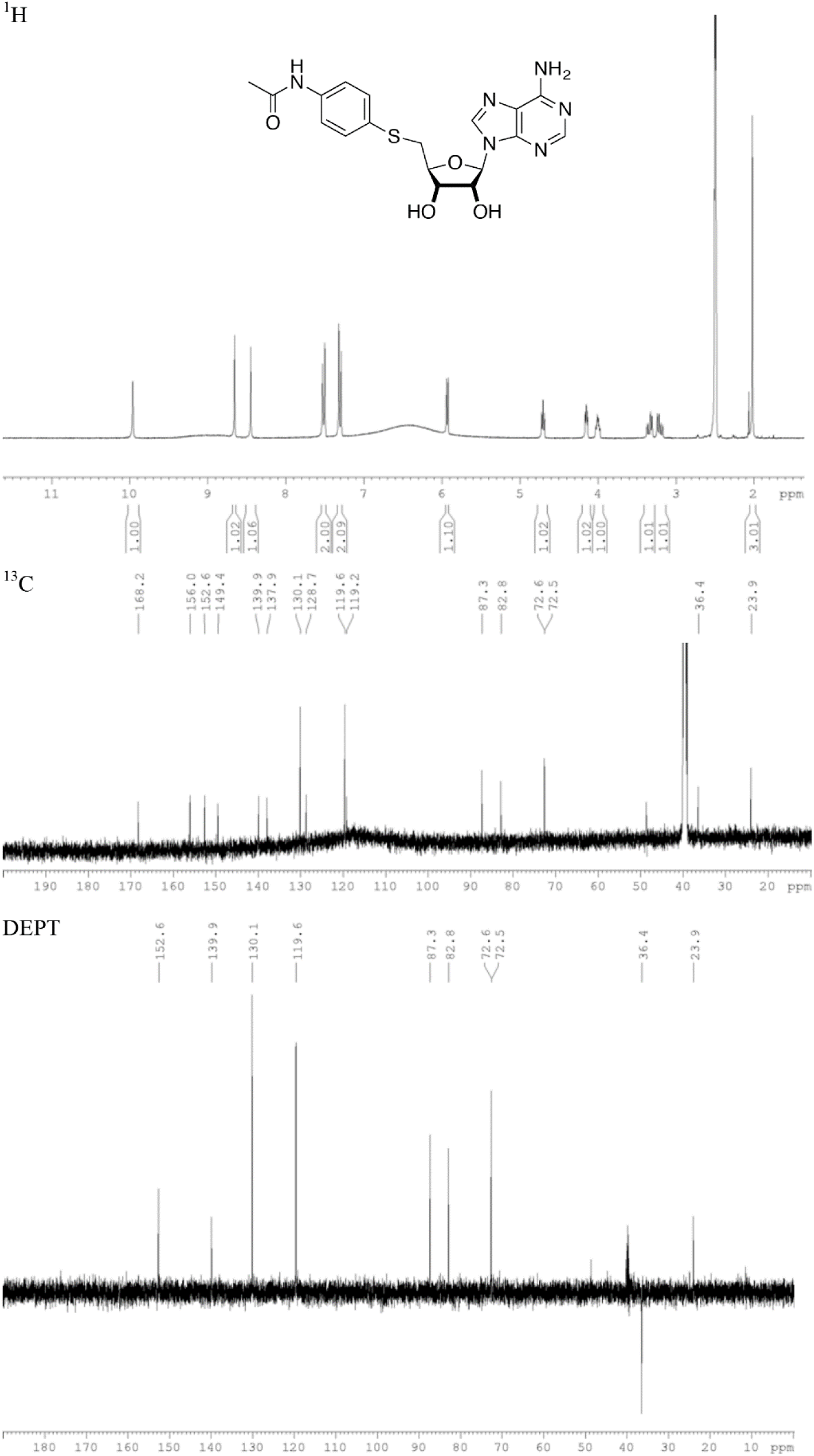
^1^H, ^13^C and DEPT-135 NMR spectra for derivative **10** in DMSO-d_6_ at 500 and 125 MHz

**Fig. S25.**
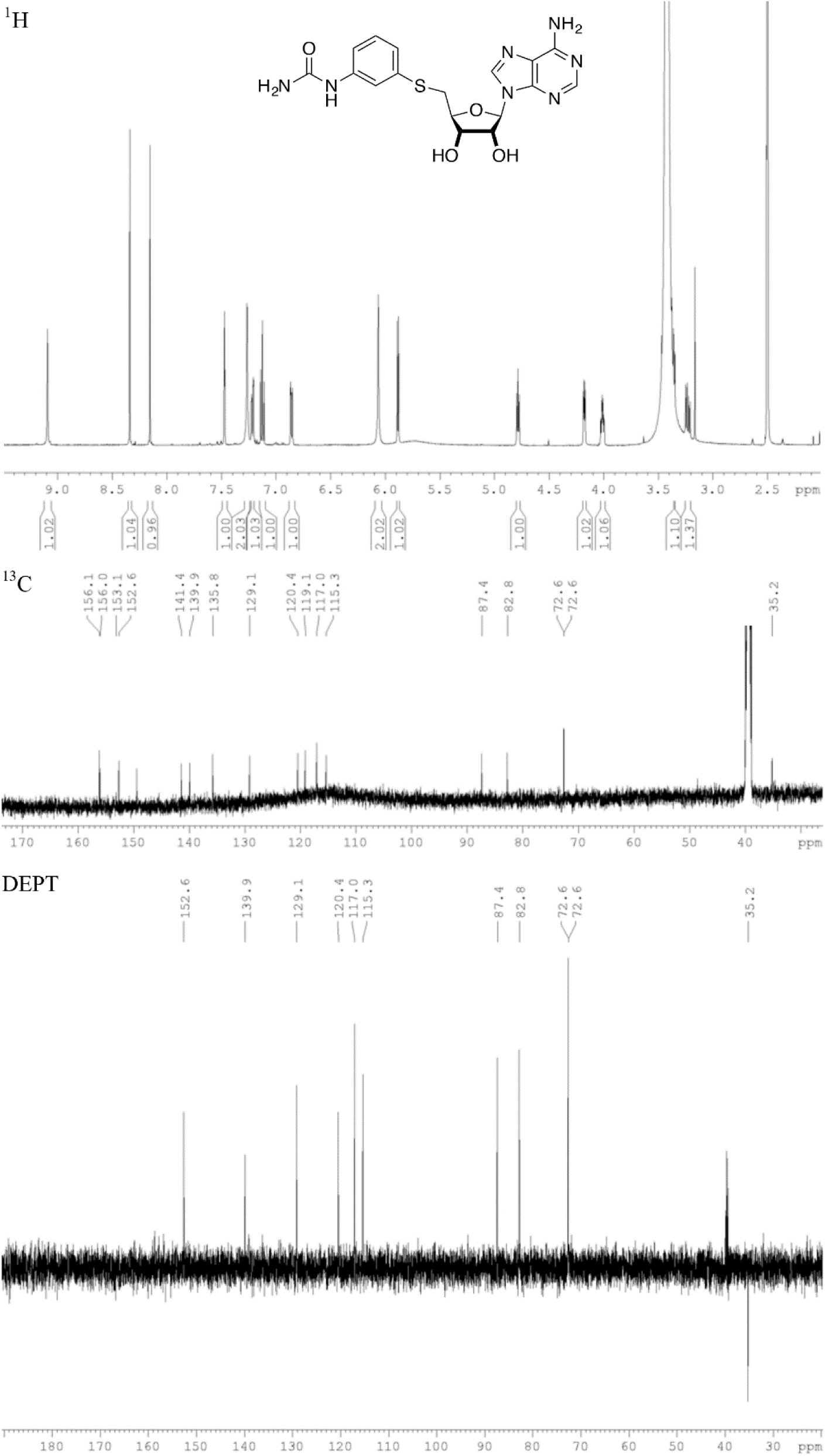
^1^H, ^13^C and DEPT-135 NMR spectra for derivative **11** in DMSO-d_6_ at 500 and 125 MHz

**Fig. S26.**
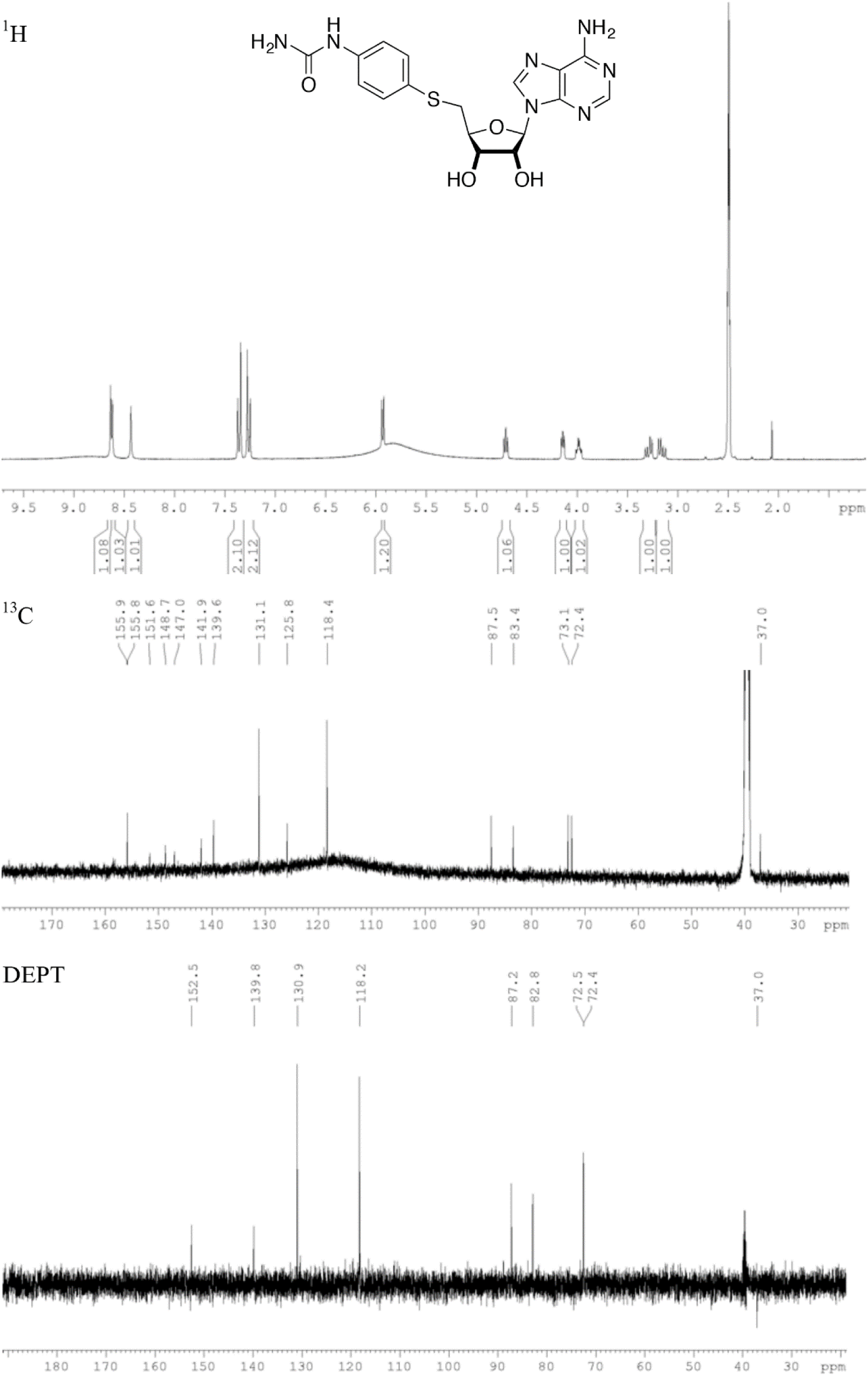
^1^H, ^13^C and DEPT-135 NMR spectra for derivative **12** in DMSO-d_6_ at 500 and 125 MHz

## DNA sequencing

Genomic DNA (2 µg) in a 96-well format was fragmented by Covaris E210 sonication for 30 seconds using a “Duty cycle” of 20% and “Intensity” of 5. The paired-end sequencing library was prepared following the BC Cancer Agency’s Genome Sciences Centre 96-well Genomic ∼350bp-450bp insert Illumina Library Construction protocol on a Biomek FX robot (Beckman-Coulter, USA). Briefly, the DNA was purified in a 96-well microtitre plate using Ampure XP SPRI beads (40-45 µL beads per 60uL DNA), and was subject to end-repair, and phosphorylation by T4 DNA polymerase, Klenow DNA Polymerase, and T4 polynucleotide kinase respectively in a single reaction, followed by cleanup using Ampure XP SPRI beads and 3’ A-tailing by Klenow fragment (3’ to 5’ exo minus). After cleanup using Ampure XP SPRI beads, picogreen quantification was performed to determine the amount of Illumina PE adapters used in the next step of adapter ligation reaction. The adapter-ligated products were purified using Ampure XP SPRI beads, then PCR-amplified with Phusion DNA Polymerase (Thermo Fisher Scientific Inc. USA) using Illumina’s PE indexed primer set, with cycle conditions: 98°C for 30 sec followed by 6 cycles of 98°C for 15 sec, 62°C for 30 sec and 72°C for 30 sec, and a final extension at 72°C for 5 min. The PCR products were purified using Ampure XP SPRI beads and checked with Caliper LabChip GX for DNA samples using the High Sensitivity Assay (PerkinElmer, Inc. USA). PCR product of the desired size range was gel purified (8% PAGE or 1.5% Metaphor agarose in an in-house custom-built robot), and the DNA quality was assessed and quantified using an Agilent DNA 1000 series II assay and Quant-iT dsDNA HS Assay Kit using Qubit fluorometer (Invitrogen), then diluted to 8nM. The final concentration was confirmed by Quant-iT dsDNA HS Assay prior to Illumina Sequencing.

